# High-Throughput Measurement of Metastable DNA Secondary Structures using Multiplexed Low-Yield Bisulfite Sequencing (MLB-seq)

**DOI:** 10.1101/2021.05.21.445174

**Authors:** Jiaming Li, Jin H. Bae, Boyan Yordanov, Michael X. Wang, Andrew Phillips, David Yu Zhang

**Author notes:** Authors contributed equally (Jiaming Li, Jin H. Bae, Boyan Yordanov). Correspondence may be addressed to DYZ or AP.

## Abstract

Predicting DNA secondary structures is critical to a broad range of applications involving single-stranded DNA (ssDNA), yet remains an open problem. Existing prediction models are limited by insufficient experimental data, due to a lack of high-throughput methods to study DNA structures, in contrast to RNA structures. Here, we present a method for profiling DNA secondary structures using multiplexed low-yield bisulfite sequencing (MLB-seq), which examines the chemical accessibility of cytosines in thousands of different oligonucleotides. By establishing a probability-based model to evaluate the consensus probability between MLB-seq data and structures proposed using NUPACK software, we identified the secondary structures of individual ssDNA molecules and estimated the distribution of multiple secondary structures in solution. We studied the structures of 1,057 human genome subsequences and experimentally confirmed that 84% adopted two or more structures. MLB-seq thus enables high-throughput ssDNA structure profiling and will benefit the design of probes, primers, aptamers, and genetic regulators.

## INTRODUCTION

The secondary structure of single-stranded DNA (ssDNA) largely determines the chemical accessibility of different nucleotides and regulates many biological processes such as transcriptional efficiency [1], splicing patterns [2], degradation rates [3], mutation rates [4], and ribozyme catalytic activity [2]. Also, many biomedical technologies now demand a reliable prediction of ssDNA secondary structures, including DNA aptamer library design [5, 6], genetic circuit development [7–9], gene therapy [10], and hybridization probe design [11]. Recently, new species and classes of ssDNA viruses [12, 13] and circular DNA [14, 15] have been discovered by shotgun sequencing analysis of microbiomes. The secondary structure of DNA molecules can be a critical link between the identified DNA sequences and the mechanisms underpinning their biological function.

Software tools for predicting secondary structures of nucleic acids from sequence have been in wide use for 30 years [16–21], but the accuracy of these tools is limited, particularly for longer biological sequences [22]. A significant bottleneck for the development of better prediction tools is the small size of available experimental structure-sequence correlation data for fitting the thermodynamic model parameters, which are mostly obtained from low-throughput melting temperature experiments.

Despite its significance, there has not been a high-throughput method for the systematic study of ssDNA structures, in contrast to RNA structure study. Methods such as SHAPE-Map [23–26] and DMS-seq [27, 28] successfully adopt next generation sequencing (NGS) to dramatically improve their throughputs when probing RNA structures, but they rely on chemical modification of RNA bases and reverse transcriptase, which are not directly applicable to DNA. However, the underlying principles of the RNA structure study methods -- which first mutates or truncates a fraction of nucleotides with a strong bias in favor of unpaired bases, reads them out through NGS, and then identifies the unpaired bases by their high reaction yields -- could be implemented for DNA structure study if structure-biased base mutations can be introduced in ssDNA. We considered bisulfite conversion since it has been previously used for transforming deoxycytosines to deoxyuracils and was reported to be structure-biased (Fig. 1a) [29, 30].

**Figure 1:**
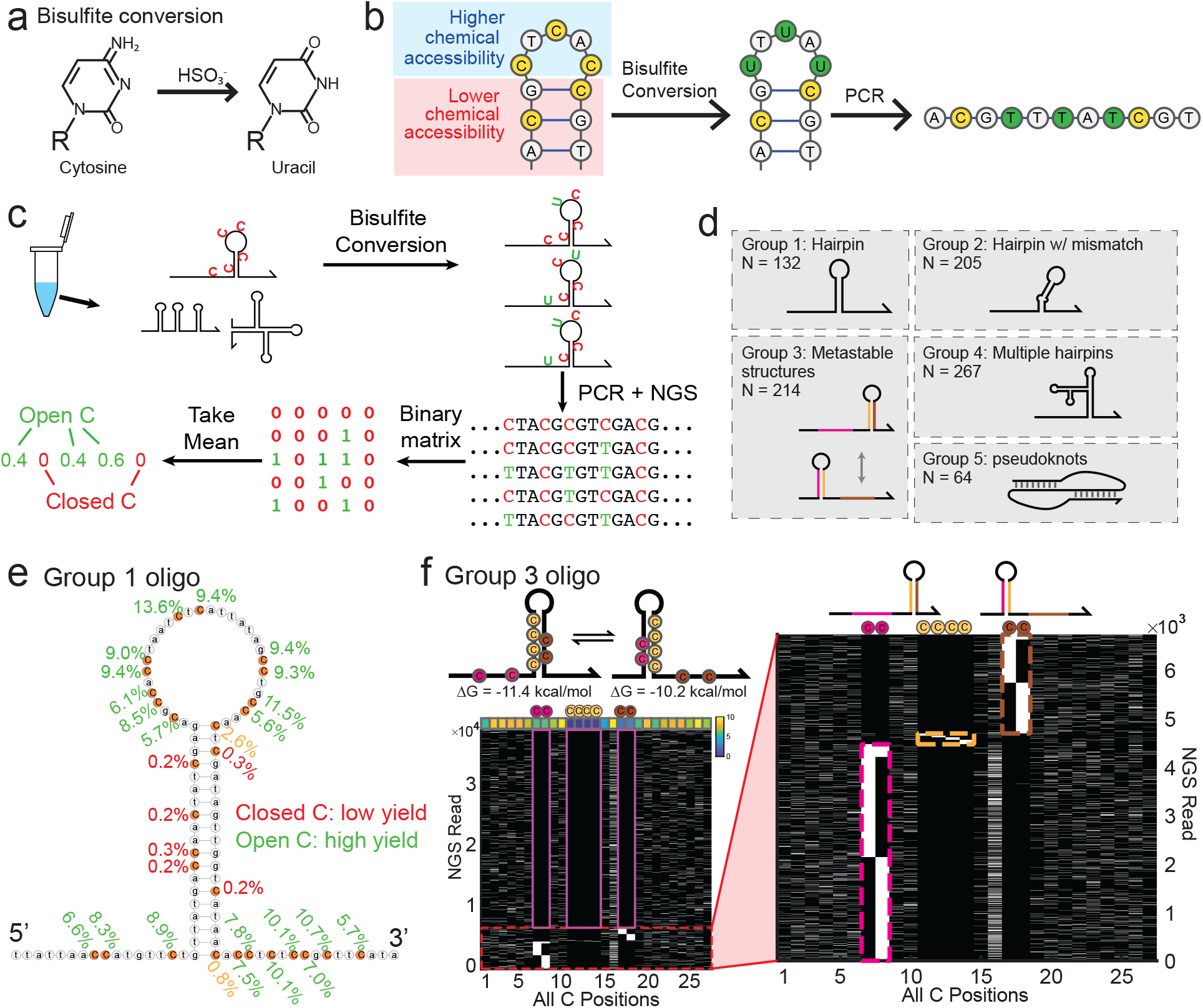
The concept, workflow, and proof of feasibility of MLB-seq. (a) Bisulfite conversion. R stands for the ribose or deoxyribose in RNA/DNA. (b) DNA secondary structure information is reflected as DNA sequence changes through the course of bisulfite conversion. C nucleotides on the hairpin loop are converted into U with higher probability than C nucleotides on the hairpin stem. (c) Low-yield bisulfite sequencing (MLB-seq) workflow overview. (d) 882 rationally designed oligos divided into 5 structural groups. (e) The conversion yields (CY) of all C positions on a representative group 1 oligo. Group 1 oligos were designed to have a stable single-hairpin structure, and all C nucleotides were colored orange for easier viewing. The CY of each C was marked by its side, green for open and red/yellow for closed. The two C nucleotides marked by yellow were at the ends of the duplex stem. (f) Analysis of a representative group 3 oligo, with two competing metastable hairpin secondary structures. The bottom left panel shows the conversion patterns of all reads mapped to this oligo. Each row represents an individual NGS read and each column represents a different C nucleotide in the DNA sequence ordered from 5’ to 3’. Black denotes unconverted (readout as C) and white denotes converted (readout as T). The upper row of color-mapped grids shows the overall conversion yield of each C position. The purple boxes mark the eight C positions shown on the oligo sketch. The oligos are not drawn to scale.

Here, we present multiplexed low-yield bisulfite sequencing (MLB-seq), a massively parallel method for chemically probing the secondary structures of many single-stranded DNA species. Importantly, multiple co-existing structures of the same DNA species can be identified through this approach, allowing biophysical characterization of the ensemble of molecules in solution. In MLB-seq, a heterogeneous mixture of multiple DNA species is reacted with a dilute solution of sodium bisulfite to convert a small fraction of cytosine (C) nucleotides to uracils (Fig. 1c). DNA nucleotides base-paired in hairpin stems have significantly lower yield of conversion than unpaired DNA nucleotides. Because all C nucleotides in a single NGS read correspond to haplotype-phased information about a single molecule of DNA, the NGS data reflects a high-throughput single-molecule assay of DNA secondary structures. We developed a probability-based computational model to calculate the probability of any particular NGS read being generated by a proposed secondary structure. By comparing the probabilities of two or more different secondary structures, we can determine the most likely secondary structures for many of our NGS reads and infer a solution distribution of multiple coexisting structures for each target DNA species. Applying MLB-seq to 1,057 subsequences of the human genome, each 100 nucleotides (nt) long, we found that 84% of them adopted 2 or more secondary structures in solution.

## RESULTS

### Bisulfite Conversion Yield Depends on Secondary Structure

Bisulfite conversion is a chemical reaction in which unmethylated cytosine (C) nucleotides are converted into uracil (U) nucleotides when the nucleic acid is treated with sodium bisulfite solution (Fig. 1a). The efficiency of the bisulfite conversion reaction is lower when the C nucleotide is base paired to a complementary guanine (G) nucleotide, as compared to when the C nucleotide is in a single-stranded state. Base-paired C’s are henceforth referred to as “closed”, and unpaired C’s are referred to as “open”. To overcome this efficiency bias for epigenetic profiling purposes, typical bisulfite conversion reactions use very high sodium bisulfite concentrations (e.g. 5 M) and multiple thermal annealing steps to ensure near 100% conversion yield (CY) for all cytosines despite the presence of secondary structures [31–33]. In our case, the bias in CY becomes a promising feature that can be leveraged to provide information regarding the structural state of each C nucleotide in a DNA molecule without involving any enzyme or other macromolecules (Fig. 1b).

To systematically characterize and validate the MLB-seq method, we purchased a pool of 1,939 DNA oligo species from Twist Bioscience for experimental studies. Among these DNA oligo species, 882 were rationally designed to have well-defined secondary structures. These rationally designed oligos were further divided into 5 groups, based on the designed secondary structure (Fig. 1d). The other 1,057 oligos were biological subsequences selected from 212 human genes. The entire DNA oligo pool simultaneously underwent the MLB-seq library preparation procedure (Fig. 1c, see also MATERIALS AND METHODS). The NGS FASTQ reads were first aligned to the 1,939 DNA oligo sequences, and then each oligo target was individually analyzed. We assumed that the rationally designed oligos from Groups 1, 2, and 4 folded stably at their designed structures and used them as positive controls. This assumption was later verified.

We first analyzed the Group 1 oligos, DNA molecules designed and predicted to have a single, highly stable hairpin structure (Fig. 1e). From the analysis of C nucleotides in these oligos, a clear pattern emerged in which open C nucleotides exhibited significantly higher CYs than closed C nucleotides (Fig. 2a, Fig. S5), consistent with our expectations. The very low but non-zero (~0.4%) CY of C nucleotides in hairpin stems might be an artifact of DNA sequencing. It is well known that sequencing-by-synthesis possesses an intrinsic error rate of between 0.2% and 1%, which is consistent with the observed CY of closed C nucleotides.

**Figure 2:**
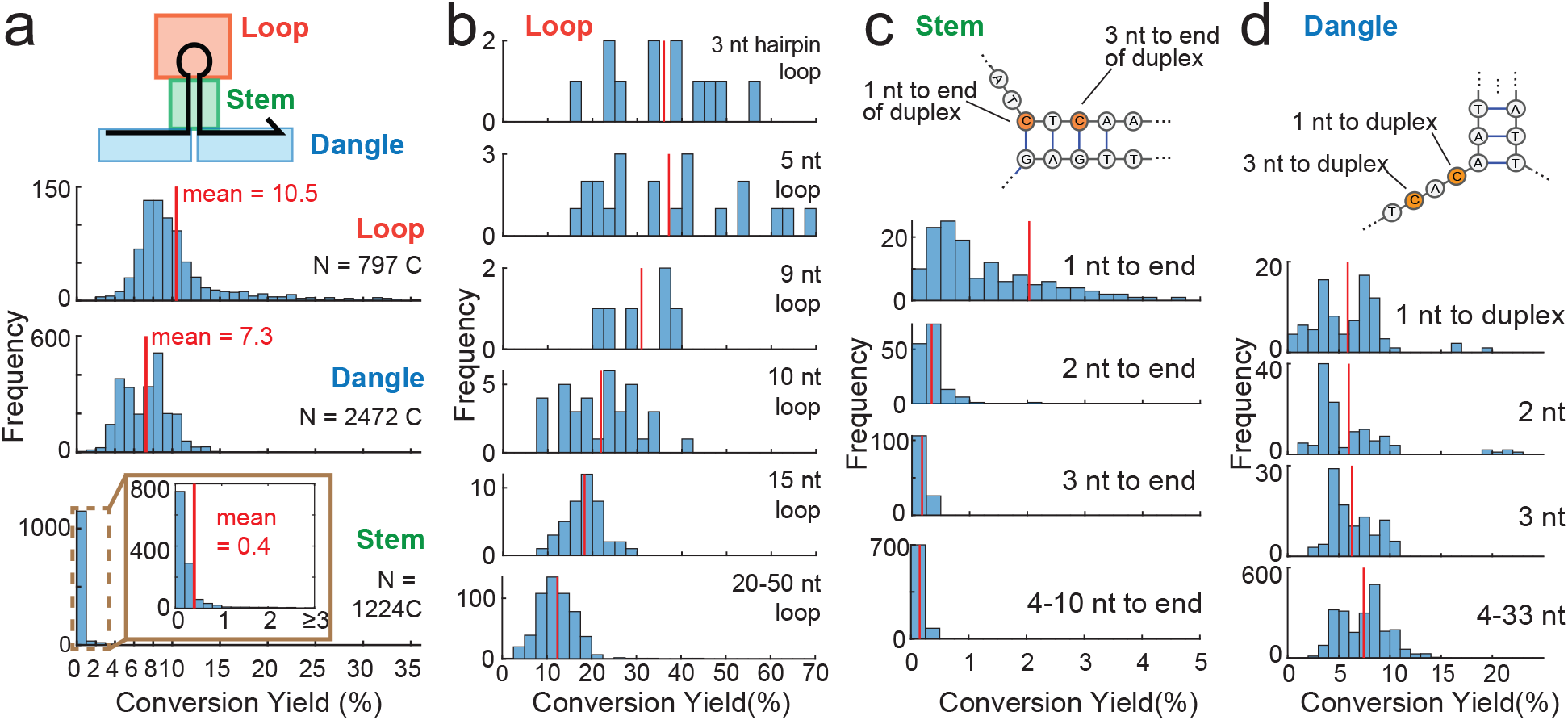
Distribution of bisulfite CY for C nucleotides summarized from 132 Group 1 oligos. (a) C nucleotides in hairpin loops and dangles have higher CY than C nucleotides in hairpin stems. (b) Smaller hairpin loops that are topologically constrained are observed to exhibit higher CY values. (c) The first nucleotide in a hairpin stem typically has higher CY compared to other hairpin stem nucleotides, possibly due to base pair breathing. (d) The position of C nucleotides in dangles does not appear to have a significant effect on CY.

We also identified two interesting phenomena in addition to confirming the impact of base pairing on CY: First, C nucleotides in smaller hairpin loops, which were expected to be more topologically constrained, exhibited 3-fold higher CY than those in larger hairpin loops and dangles (Fig. 2b). One possible explanation for this behavior is that the C nucleotides in larger hairpin loops and dangles can transiently base pair with G nucleotides on other DNA species in solution, but C nucleotides in small hairpin loops are physically incapable of base pairing. Second, the C nucleotides in a hairpin stem closest to the end of the stem exhibited higher CY than C nucleotides near the middle of a hairpin stem (Fig. 2c), suggesting that base pair breathing may have resulted in higher CY. In contrast, no such position dependence was observed for the distance of open C nucleotides to the hairpin stem (Fig. 2d). This is consistent with similar observations in RNA structure probing [33, 34]. We carried out the same analysis for Group 2 and Group 4 DNA oligos and obtained consistent results (Fig. S8). We thoroughly analyzed and compared data from the other 5 NGS libraries and found good reproducibility and consistency between libraries (Fig. S4, S5).

Group 3 oligo sequences were designed to adopt multiple metastable secondary structures (Fig. 1f). We used them to demonstrate that MLB-seq can identify distinct subpopulations of molecular states. In Fig. 1f, the conversion of the C nucleotides in the 3 pairing regions (as marked by the magenta, yellow, and brown dashed line boxes) were nearly mutually exclusive, corresponding to the two prospective single-hairpin structures. The reads in the magenta box with C nucleotides converted in the left pairing region corresponded to molecules folded into the structure shown on the left, and the reads in the brown box with C nucleotides converted in the right pairing region corresponded to the structure shown on the right. The four yellow C nucleotides in the middle paring region had very few conversions because they were closed in both structures. These mutually exclusive conversions demonstrated the capability of MLB-seq identifying metastable secondary structures at a single molecule level and measuring the relative concentrations of the co-existing structures.

### Probability model

We aimed to use MLB-seq to identify structural information of ssDNA. One approach is to use CY as a direct measurement of the open probability (OP) of each C nucleotide. We compared the measured CY with OP predicted by NUPACK and found good correlation between the two for oligos in groups 1 to 4 (R = 0.74, 0.84, 0.82, and 0.81, Fig. S9). This confirmed CY to be a good measurement of OP for these oligos. However, for the biological sequences the correlation between CY and OP became weaker (R = 0.56). We hypothesized that this was because the NUPACK predicted OP is calculated from the ideal equilibrium states using thermodynamic model, but in solution oligos might be in other metastable structural states due to fluctuating folding kinetics, resulting in different CY. This identified the potential for MLB-seq to provide a measurement of the metastable structural states of biological oligos, which are more difficult to predict.

To validate our hypothesis and demonstrate the forementioned capability of MLB-seq, we developed a probability-based computational model that analyzed each NGS read from MLB-seq and indicated the most likely structure of the molecule generating this NGS read (Fig. 3). To do so, we first made a simplifying assumption that the probability of a C nucleotide conversion is dependent only on the base pairing state of this nucleotide, and otherwise independently and uniformly distributed. Then, using Bayes’s rule, we calculated the posterior probabilities of a C nucleotide being open or closed during bisulfite conversion, given it is observed converted or not in the MLB-seq readout. See MATERIALS AND METHODS and Supplementary Text for calculation steps of these probabilities.

**Figure 3:**
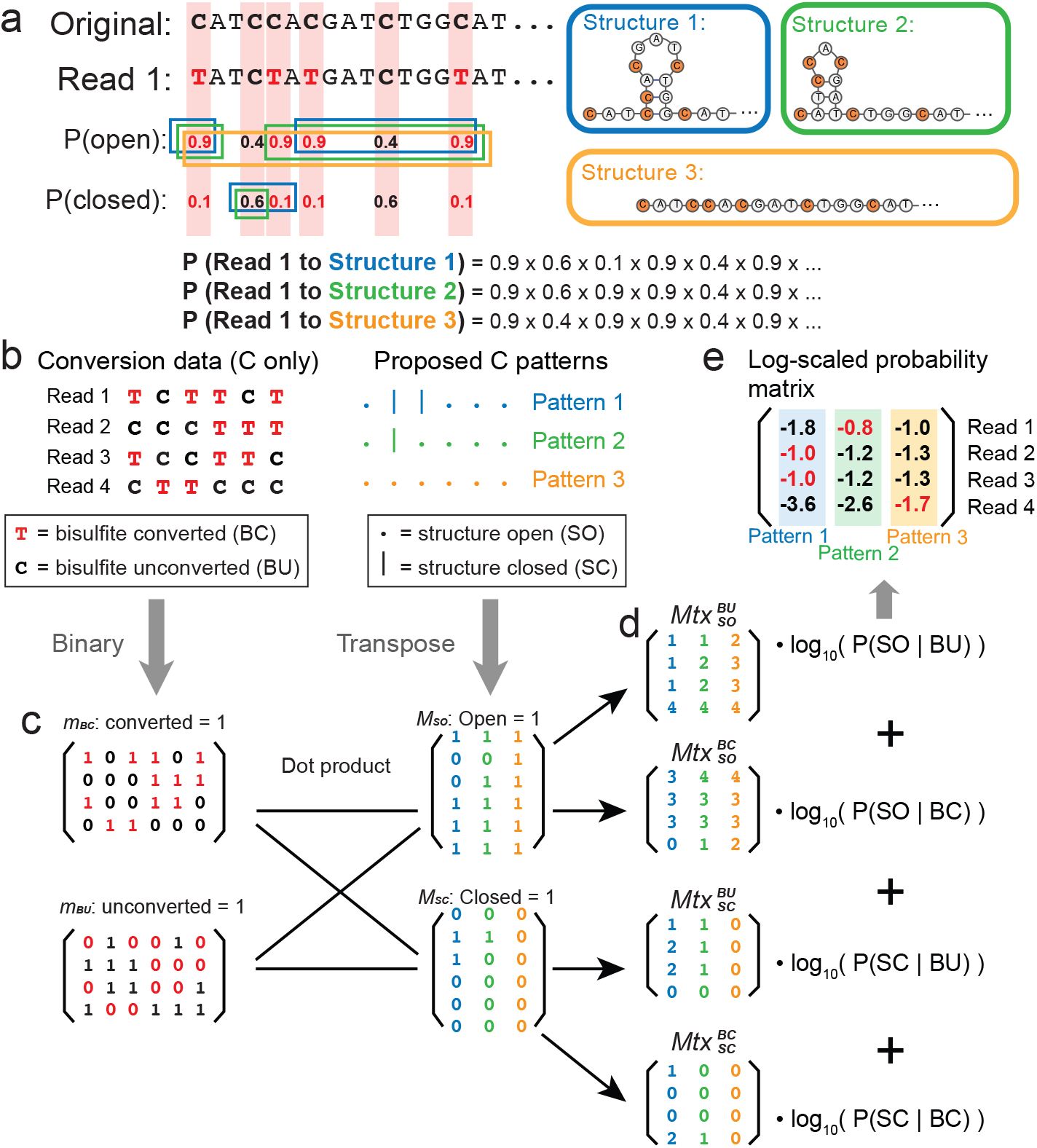
Conceptual model demonstrating the framework for calculating probabilities of secondary structures from observed NGS reads. (a) Illustration of probability calculation based on an observed NGS read. All probabilities shown in the figure are simplified for demonstration and are not the actual values used throughout the analysis. (b) Conversion state data from NGS reads and predicted structure open/closed patterns serve as the two input matrices. (c) Two pairs of complementary binary matrices are generated, and the pairwise dot products are calculated. (d) These four matrices from dot product measure the number of C nucleotides in the four orthogonal conditions: being unconverted and open, being converted and open, being unconverted and closed, and being converted and closed. The numbers at the same position of the four matrices come from the same read and structure combination. For example, the element on the *i^th^* row and *j^th^* column of matrix 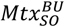 represents the number of C nucleotides in the *i^th^* read being both unconverted in MLB-seq and open in the *j^th^* candidate structure. *P*(*SO|BC*), *P*(*SC|BC*), *P*(*SC|BU*), and *P*(*SO|BU*) are the posterior probabilities of one C nucleotide being open or closed given it observed converted or not. Here P stands for posterior probability, *SO* and *SC* stands for structure open and closed, and *BC* and *BU* stand for bisulfite converted and unconverted. (e) The sum of the matrices is the output: the log-scaled probability of each read corresponding to each proposed structure.

For a target oligo whose structure is uncertain, we first pre-selected a pool of candidate structures (please see MATERIAL AND METHODS). We then aligned an MLB-seq read to its original sequence and obtain the conversion states of each C. For one candidate structure, we multiplied the posterior probabilities of all C nucleotides being open or closed according to the structure, given their observed conversion states from the NGS read. We defined the multiplication result as the probability of this NGS read corresponding to the candidate structure. In this way, we calculated the probabilities of every individual NGS read corresponding to different candidate structures (Fig. 3a), and we were then able to map all the reads individually to the structures with the highest probability.

To facilitate higher computing efficiency with our high-throughput NGS data, we utilized linear transformation via matrix calculation. For each target oligo, the model started with two symbolic matrices as inputs: (1) the conversion data and (2) the open and closed patterns of all C positions from the candidate structures (Fig. 3b). We obtained the conversion data exclusively from MLB-seq experimental readouts, and we used the NUPACK software [18, 19] to generate the candidate structures. The two inputs were each processed into a pair of complimentary binary matrices (Fig. 3c): a converted matrix and an unconverted matrix from the conversion data (denoted as *m_BC_* and *m_BU_*), as well as an open matrix and a closed matrix from the structural patterns (denoted as *M_SO_* and *M_SC_*). We then applied dot products to combine conversion data with structural patterns and acquired four matrices (Fig. 3d):

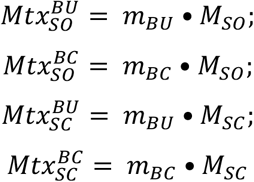

These four matrices measured the number of C nucleotides in the four orthogonal conditions: being unconverted and open, being converted and open, being unconverted and closed, and being converted and closed. The number of C nucleotides in each conversion and structure combination equals the exponent by which we should multiply the posterior probability of this condition. By taking the logarithm of the posterior, the exponentiation calculation was transformed into simple multiplication. By multiplying the four matrices by the four log-scaled probabilities respectively and summing them, we obtained the output matrix: the log-scaled probabilities of every individual NGS read corresponding to each candidate structure (Fig.3e). The highest (least negative) probability in each row indicateed the most likely structure of the molecule generating this NGS.

### Model validation with arbitrarily designed oligos

We examined the model with MLB-seq data of the positive control oligos from Groups 1, 2, and 4. For each oligo, we fed two candidate structures into the model: the designed minimum-free-energy (MFE) structure as structure 1, the positive reference, and a thermodynamically unfavored structure as structure 2, the negative reference. Details on the generation of these thermodynamically unfavored structures can be found in Supplementary Text.

Fig. 4a shows the model output of an oligo target from group 1, with the log-scaled probabilities of each read corresponding to each structure plotted as a heatmap. The two columns represent structure 1 (Fig. 4b) and structure 2 (Fig. 4c), and the rows represent all the NGS reads from this oligo. The color of each grid represents the log-scaled probability of this molecule (NGS read) folded at the corresponding structure. The heatmap has been sorted in the vertical axis for easier viewing. About 7,000 reads were consistent with both structures, as clustered on the upper half of the heatmap, and we defined them as ‘non-specific’ reads. This is understandable since the overall reaction yield was low and there might be a number of molecules without sufficient conversions to specifically distinguish between the two structures. All other reads that distinctively support one structure were defined as ‘non-ambiguous’ reads. Details on how we classified the non-specific and non-ambiguous reads in consistency are included in Supplementary Text.

**Figure 4:**
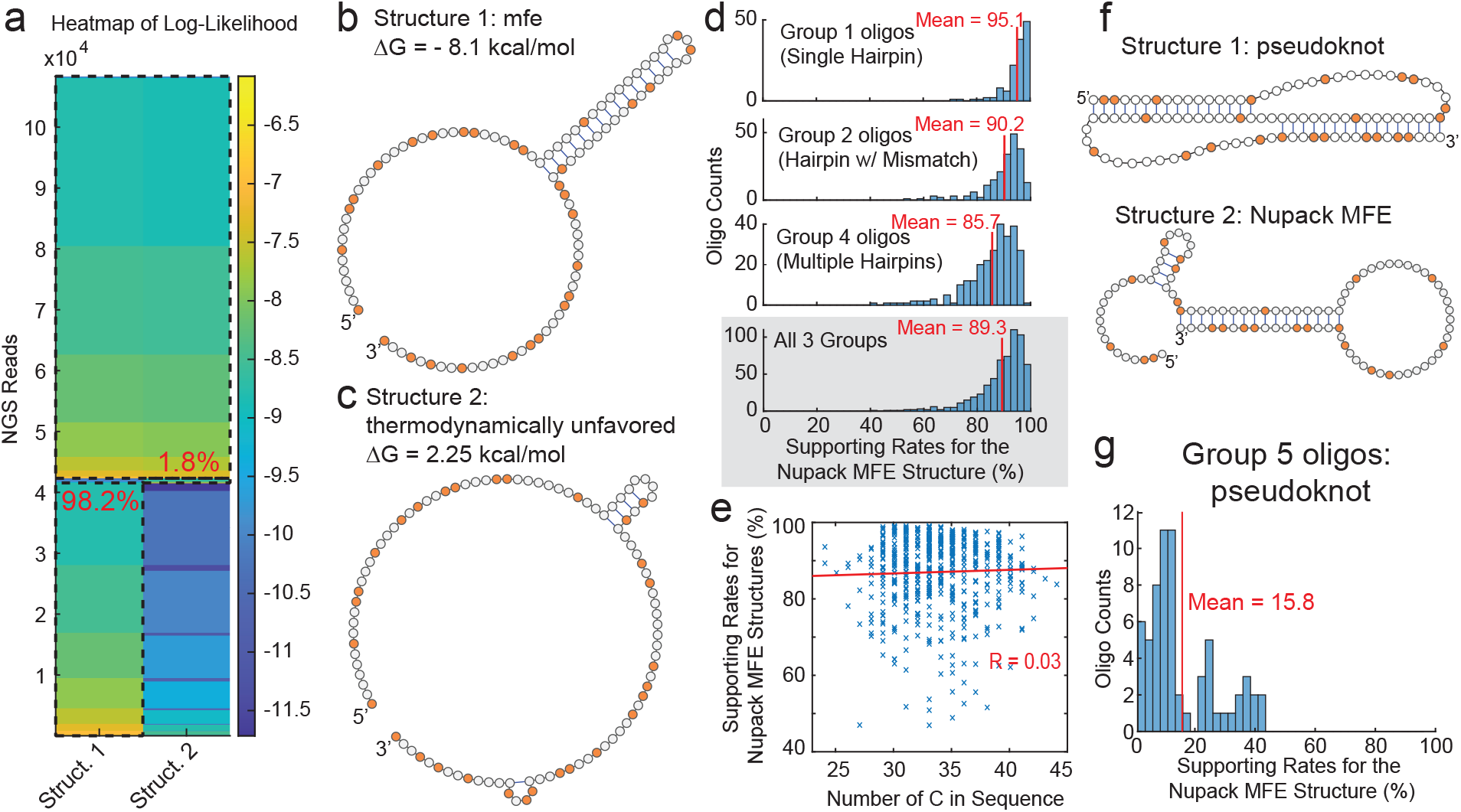
Validation of the model using MLB-seq data from arbitrarily designed oligos. (a) Heatmap of log-scaled probabilities of one oligo target. The black dashed-line boxes classify reads supporting structure 1, reads supporting structure 2, and the nonspecific reads that are consistent with both structures. The two percentage values colored in red are supporting rates for the two structures. (b) Structure 1, the NUPACK MFE structure of the oligo. Used as the positive reference. (c) Structure 2, a structure generated to be significantly less thermodynamically favorable. Used as the negative reference. (d) Summary of the supporting rates for the NUPACK MFE structures from oligos in Groups 1, 2, and 4. (e) Summary of the supporting rates for the NUPACK MFE structures against the number of C nucleotides in sequence. Summarized from all oligos in Groups 1, 2, and 4. (f) Expected pseudoknot structure (structure 1, positive reference) and NUPACK predicted MFE structure (structure 2, negative reference) from an example oligo in Group 5. (g) Summary of the supporting rates for the NUPACK MFE structures from Group 5 oligos.

We calculated the supporting rate for each structure by dividing the number of reads distinctively supporting this structure by the number of all non-ambiguous reads. For this target shown, the MFE structure was significantly supported by more than 4,000 reads, resulting in a supporting rate of 98.2%. We considered this as the true positive rate. The remaining 1.8% of reads supporting structure 2 may be the result of sequencing errors, as described earlier. Fig. 4d summarizes the supporting rates of all designed oligos in Groups 1, 2, and 4; almost all of these exhibited high true positive rates. We also ensured that the model performed robustly regardless of the varied number of C nucleotides in the sequences (Fig. 4e, S10c, d). Meanwhile, the fact that most of the positive control oligos generated a majority of non-ambiguous reads with high true positive rates verified our assumption that these oligos folded at their predicted structures stably in solution during MLB-seq and could be used as positive control (Fig. S11e).

To test whether MLB-seq and our probability model can contradict incorrect structures, we considered the Group 5 oligos, that were all arbitrarily designed to have a pseudoknot secondary structure. A pseudoknot structure occurs when nucleotides in the loop of a hairpin participate in hybridization to a set of nucleotides outside of the loop. Due to the technical details of the Nussinov algorithm for base-pair-maximum structure prediction, DNA secondary structures with pseudoknots are hard to predict. Thus, the NUPACK-predicted MFE structures for Group 5 oligos did not have pseudoknots and were expected to be incorrect. For all Group 5 oligos, we compared the NUPACK-predicted MFE structure (Structure 1) with the expected pseudoknot structure (Structure 2, Fig. 4f), and observed significantly lower supporting rates for the NUPACK-predicted MFE structures (i.e., high true negative rate) than for Groups 1, 2, and 4 (Fig. 4g). To summarize, we observed high true positive and true negative rates, proving the good performance of the model coupled with MLB-seq. Comprehensive benchmarking of the model can be found in Supplementary Text.

### MLB-seq results on biological subsequences

We next analyzed the MLB-seq results on 1,057 human DNA subsequences from 212 human genes. The selection process, chromosome locations, exon/intron status, GC content, and NUPACK predicted MFE structures of these subsequences are summarized in the Supplement Text. Due to technical limitations of NGS library preparation, 33 of these 1,057 sequences (3.1%) had too few NGS reads for reliable statistical analysis and were discarded.

We first constructed a pool of potential secondary structures for each DNA sequence using the NUPACK suboptimal function [18, 19]. Because a particular sequence could form many structures that were highly similar with few base pairing changes (e.g. base pair breathing at the end of hairpin stems), these structures did not generally represent distinct metastable structural states that the molecules adopted in solution. Thus, we clustered these potential structures by a hard-set Hamming distance threshold and selected the structure with the lowest folding energy in every cluster as a candidate structure (Fig. 5a).

**Figure 5:**
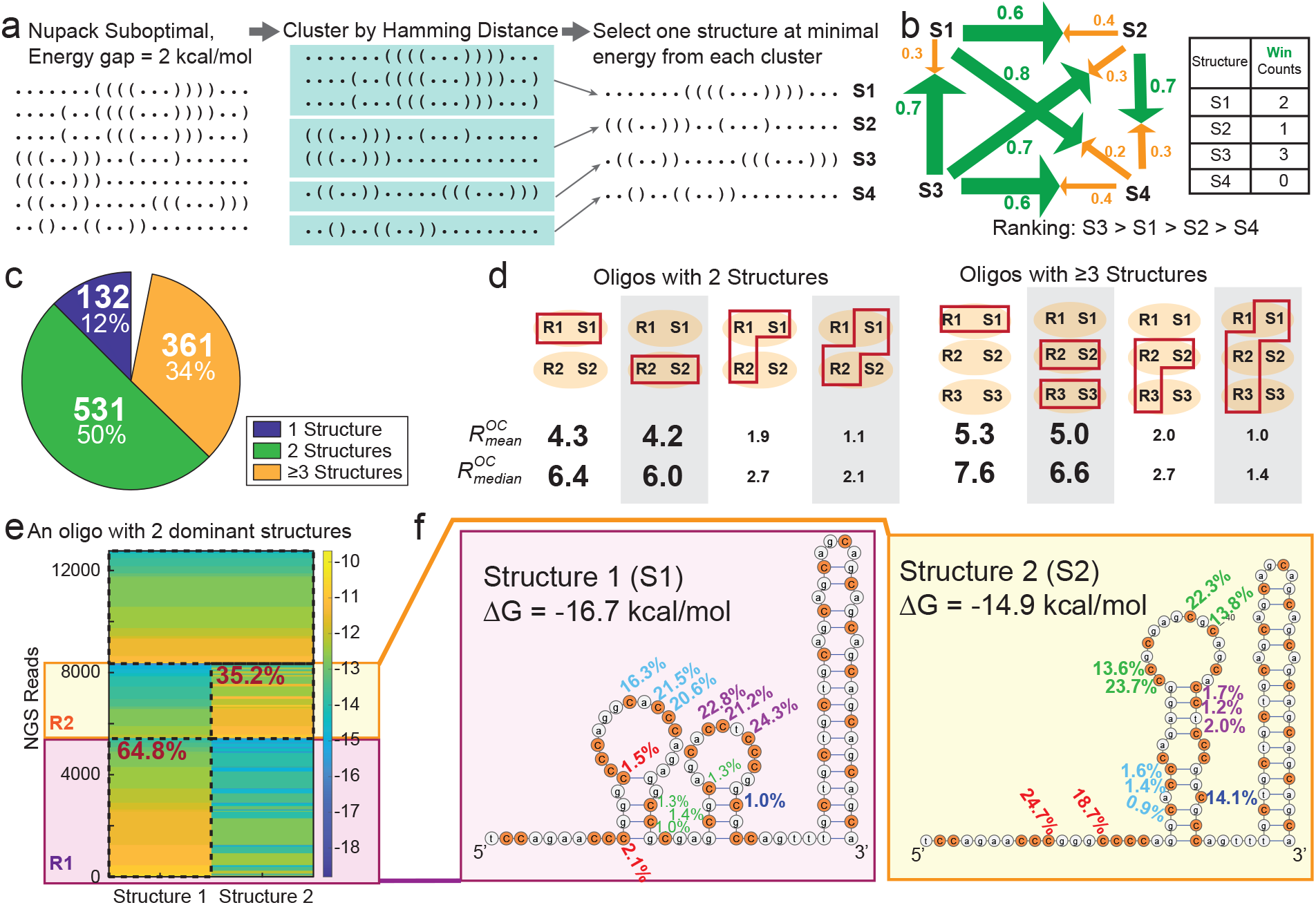
MLB-seq and the probability model shows that most biological DNA sequences adopt two or more co-existing structures. (a) Toy model demonstrating how we generate the candidate structures for each oligo target. (b) Pairwise comparison and Condorcet winning strategy that determines the dominant secondary structures for final output. Here we use 4 candidate structures (S1, S2, S3, and S4) as a toy model. Supporting rates are marked by the arrows. The number of winning matchups is tallied for each structure. (c) Distribution of the number of co-existing secondary structures observed in human DNA subsequences via MLB-seq. The remaining 33 sequences (out of 1,057) had too few NGS reads for confident secondary structure determination. (d) Verification of multiple co-existing structures. 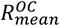 and 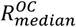 represents the mean and median relative ratio CY for C nucleotides predicted to be open vs. those predicted to be closed. When the structure is supported by MLB-seq NGS reads, we expect high *R^OC^* values. R1 represents the NGS read subset that distinctively supports structure 1 (S1), and so does R2 and R3, S2 and S3, respectively. The red boxes mark the NGS read subsets used for CY calculation and the structure that defines the open and closed states of C nucleotides. (e) Model output of one biological DNA subsequence with strong evidence for co-existence of 2 structures. (f) The two co-existing structures from the target in panel (c). The CY calculated using only R1 or R2 are marked next to corresponding C nucleotides, with the same colors representing the same loci.

For each sequence, we compared all candidate structures in the model pairwise and adopted a Condorcet winning strategy to select the dominant one, two, or three structures as the final output (Fig. 5b). Their supporting rates indicated the fraction of DNA molecules that adopted these structures in solution (See MATERIALS AND METHODS for details). We found that 84% of the 1,057 human DNA subsequences adopt 2 or more co-existing DNA secondary structures in solution (Fig. 5c).

We verified our conclusion with quantitative metrics. Because open C nucleotides have higher CY than closed C nucleotides (Fig. 2a), the quality of a secondary structure prediction from a particular set of NGS reads can be quantitated as *R^OC^*, the ratio of the CY for the predicted open C nucleotides divided by the CY of the predicted closed C nucleotides. For our rationally designed DNA sequences, the value of *R^OC^* was roughly 20 (8% vs. 0.4%). In contrast, a random secondary structure with no grounding would presumably result in *R^OC^* ~ 1. The higher the value of *R^OC^*, the more confidence we have in the correctness of the structures.

Consequently, the MLB-seq data for DNA sequences that adopted two distinct co-existing secondary structures can be divided into two subsets of reads, R1 and R2, with very different *R^OC^* values when compared against the two structures S1 or S2. For oligos with 2 co-existing structures, the R1 reads produced high *R^OC^* values when associated to S1, but when the entire R1 and R2 datasets were combined, there was low *R^OC^* value to S1 (Fig. 5d). Fig. 5e shows an example with strong evidence for co-existence of two structures as demonstration. The CY of all C nucleotides were calculated using only R1 or R2 reads. All C nucleotides in different open/closed states in the two structures generated distinctive high CY (>13%) when open and low CY (≤2.1%) when closed, consistent with our previous observation.

## DISCUSSION

We established MLB-seq by integrating the structure-biased bisulfite conversion of C nucleotides with NGS and used it to query the secondary structures of ssDNA targets with high throughput. By studying the reaction yields of 604 rationally designed positive control oligos, we developed a statistical model that calculated the probability of an individual MLB-seq molecular readout originating from any proposed secondary structure. In this way, we were able to observe the distribution of co-existing secondary structures of each DNA species that went through MLB-seq. We studied 1,057 biological subsequences and found 84% of them had multiple co-existing structures in solution.

MLB-seq probes only C nucleotides. This adapts well to the nucleotide-independence assumption that enables the probability model, since C nucleotides are usually separated by other non-C nucleotides, so the conversions are more independent. Other existing RNA structure probing methods, involving specific reverse transcriptase to generate the mutations from structures, usually result in stronger interference between adjacent or nearby nucleotides due to the nature of polymerase. That being said, our probability model could be readily generalized to RNA structure probing methods as long as the independence assumption holds. On the other hand, this is a primary limitation of MLB-seq, making it not applicable to A/T-rich DNA sequences. A fraction of the NGS reads do not provide sufficient information to distinguish between two proposed secondary structures. Similar structures of one DNA species with just a few bases at different open/closed states might not be distinguishable. Therefore, a future research focus will be to investigate other base modification methods, e.g., oxidation, that target other nucleotides and can be paralleled with MLB-seq to generate denser information and higher sensitivity.

Our pipeline of generating the pool of candidate structures and feeding them into the model to select the correct ones belongs to a class of methods termed ‘sample and select’ [36]. These methods share a common difficulty: it is hard to guarantee a good pre-generated ‘sample’. In our specific case, the NUPACK suboptimal prediction itself is not completely accurate [37]; the energy gap of 2 kcal/mol does not guarantee sufficient suboptimal structures for all target oligos. In theory, a larger energy gap for the suboptimal prediction might generate samples more thoroughly. However, in practice when we increased the energy gap above 2 kcal/mol, the NUPACK suboptimal function ran significantly slower (> 1 hour) on some oligo targets and generated many structures with just a few bases of difference, which was inefficient and ill-suited to our high throughput data. Further work is needed to explore the possibility of generating candidate structures without involving thermodynamic calculation, similar to how we generated the thermodynamically unfavored structures, to generate larger structure pools at lower computational cost.

From our MLB-seq NGS data, we observed that C nucleotides in small, topologically constrained hairpins have higher CY than those in dangles and large loops, by a factor of 3 to 5 (Fig. 2a, b). For model simplicity and to maintain single-molecule resolution, we did not consider the differences in CY for different types of open C nucleotides. In principle, a more sophisticated model that attempts to infer hairpin loop size and relative position of paired C’s to hairpin stem ends can provide more accurate and detailed information to confirm or refute proposed secondary structures. Furthermore, given the kinetic nature of bisulfite conversion bias based on C nucleotide state, it is possible that integrating data from multiple longitudinal MLB-seq timepoints can give more information to ascertain secondary structure and transitions between structures.

With accelerating advances in machine learning, complex problems such as nucleic acid secondary structure prediction from sequence could potentially be more accurately solved using neural networks, rather than rule-based biophysical models. Machine learning approaches, however, require a massive number of labeled instances to serve as training data. It has so far been difficult to obtain large and accurate secondary structure datasets, due to the low-throughput nature of X-ray crystallography and other techniques. MLB-seq leverages the secondary structure dependence on bisulfite reaction kinetics and uses NGS to generate large secondary-structure informing datasets on thousands of different DNA species. In principle, this approach could be further scaled up to millions of distinct nucleic acid species, and thus enable the development of machine learning based predictors of secondary structure.

## MATERIALS AND METHODS

### Synthesized DNA oligos

All DNA oligos used in the experiment were purchased from Twist Bioscience at stock concentration of 8 ng/μL. Their sequences are included in the supplementary material ‘LYB_Sequences.xlsx’. All were synthesized at a length of 150 nt. The first 26 nt from the 5’ end and the last 24 nt next to the 3’ end are universal primer regions that are for convenience of universal amplification of the oligos and will be trimmed off before the bisulfite conversion. The middle 100-nt long regions left are the target oligos present in bisulfite reaction, either arbitrarily designed to have specific secondary structures, or subsequences truncated from human genome.

### Preparation of synthesized oligos

The initial oligo pool was diluted 100x from the 8 ng/μL stock with TE buffer and yeast tRNA as carrier. Then a 21-cycle PCR was performed using KAPA HiFi HotStart Uracil+ ReadyMix (KAPA Biosystems) and universal primers, followed by a cleanup with 1.8x AMPure XP magnetic beads (Beckman Coulter). Next the 3’ primer regions were truncated using the BciVI restriction endonuclease (NEB), followed by 2.0x AMPure XP cleanup. Then we used Lambda exonuclease (NEB) to digest the bottom strand of dsDNA and truncated the 5’ primer region with USER enzyme (NEB). We added 5 M NaCl 20% v/v to the oligo mix and pre-annealed by heating it at 95°C for 3 minutes, followed by cooling it at the speed of −1°C/10s down to 25°C.

### Bisulfite conversion

The sodium bisulfite solution was always freshly prepared for each experiment. We first degassed the nuclease-free water with ultrasonication for 10 minutes. We dissolved 950 mg of sodium metabisulfite (Sigma-Aldrich) in 9.5 mL degassed water and then added 80 μL of 10 M NaOH (Sigma-Aldrich) and 120 μL of 250 mM Hydroquinone solution (Acros Organics, 99%). Next, we carefully added 10 M NaOH until the pH of the solution reached 5.5. Then we added water until the volume reached 10 mL and the bisulfite solution was ready to use. We mixed 6 μL of pre-annealed oligo mix and 114 μL of the sodium bisulfite solution, and incubated the mixture at 55°C for 0, 2, 4, or 6 hours. After incubation, a cleanup was performed using EZ DNA Methylation Gold Kit (Zymo Research). The only adjustment different from the kit standard protocol was that we added 5 M NaCl to the binding buffer and made it to have 0.833 M NaCl for salinity compensation. All samples were then stored at −20°C until next step.

### Adapter Ligation for NGS

We ligated Illumina P7 adapters to the 3’-ends of bisulfite converted oligos following the SPLAT method [38]. We first prepared the ss1 mix with 40 μM LB-splint1 and 60 μM LB-adap3 in 49 μL TE buffer and 1 μL of 5 M NaCl. We annealed the ss1 mix by heating it at 65°C for 1 minute and cooling it at the speed of −1°C/10s down to 25°C. Then we applied the T4 DNA Ligase (NEB) to the mix of ss1 and bisulfite converted oligos for 3’-end adaptor ligation, followed by cleanup using Zymoclean Gel DNA Recovery Kit. Next, we performed 5’-end adaptor ligation following the same procedure, except that ss2 mix (LB-splint2 and LB-adap5) replaced ss1 mix and the cleanup was by 1x AMPure XP beads purification. Then the oligos went through PCR with KAPA HiFi HotStart Uracil+ Ready Mix and NEBNext Multiplex Oligos for Illumina Set 1 (NEB #E7600) as primers, followed by twice 1.2x beads purification. We sequenced the final elution via NGS.

### Extract conversion data from NGS reads

The NGS data alignment was processed with a customized F# tool. The original 100-nt sequences of the 1,969 synthesized oligos were used as the reference template for NGS read alignment, with all C nucleotides replaced by Y (C or T). We aligned NGS reads by perfectly matching every nucleotide and neglected the minor fractions of reads with base mutations other than C>T. The aligned data was then processed and analyzed in MATLAB. We only kept the bases at identical positions as those C positions in the original template sequence from the reads and converted them into a binary matrix where 0 represents C, and 1 represents T converted from C (Fig. 1c). By calculating the mean of each column of the matrix, we got the conversion yield (CY) of each C nucleotide of this oligo.

### Calculation of conditional probabilities

From all oligos in Groups 1, 2, and 4, we counted and summed the numbers of C nucleotides that were (1) open and converted, (2) open and unconverted, (3) closed and converted, and (4) closed and unconverted, denoted as *N*_*SO*&*BC*_, *N*_*SO*&*BU*_, *N*_*SC*&*BC*_, and *N*_*SC*&*BU*_ respectively. We then calculated the posterior probabilities by:

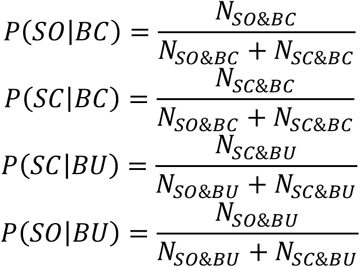

Finally, the values of the probabilities that we used throughout the model, if not specifically noted, were: *P*(*SO|BC*) = 0.93, *P*(*SC|BC*) = 0.07, *P*(*SC|BU*) = 0.45, *P*(*SO|BU*) = 0.55. All calculations above were performed in MATLAB.

### Generation of thermodynamically unfavored structures

We treated the task of generating diverse and energetically feasible (though still unfavored compared to the MFE) structures as a multi-objective optimization problem and adapted ideas from multi-objective evolutionary algorithms [39]. We focused our search on non-dominant (Pareto-optimal) structures. We started with a population of randomly generated, valid structures. Then at each iteration, we randomised the set of structures with mutations and cross-over, while we preserved only the resulting structures that were valid but also non-dominated. In the end for each oligo, we had a pool of structures with free energy ranging from MFE to slightly above 0 kcal/mol. For the model validation, we randomly selected a structure from the pool following the two restrictions below: (1) the structure had at least 10 bases at different open or closed states from the MFE structure; (2) the free energy of the structure is above −2kcal/mol.

### Analysis pipeline for biological subsequences

All analysis was carried out in MATLAB. For each biological subsequence target, we extracted the conversion data from MLB-seq NGS reads as described. We ran the ‘subopt’ function of NUPACK, with the temperature parameter set at 55 °C and the energy gap at 2 kcal/mol to generate multiple suboptimal structures. We clustered the structures using Hamming distance with a hard-set threshold of 8 nucleotides in different structural states and selected one structure from each cluster with the lowest free energy (Fig. 5a). These structures were used as candidate structures and fed into the model along with the conversion data. Next using the log-scaled probabilities acquired from the model, we made pairwise comparison between every pair of structures using all NGS reads and selected the top three structures with most wins (Fig. 5b). Finally, we compared the probabilities of all NGS reads against these three structures and the structures distinctively supported by ≥ 20 reads were defined as the dominant co-existing structures. If a target had only 1 structure generated by NUPACK within the 2 kcal/mol energy gap, we defined it the only dominant structure. If a target had 2 structures generated by NUPACK within the energy gap, we defined the structure(s) supported distinctively by ≥20 reads as the dominant structure(s).

## Acknowledgements

This work was funded by NIH grants R01HG008752 and R01HG011356 to DYZ. The authors thank Lijin Zeng for editorial assistance.

## Code Availability

NGS data alignment was processed by a tool written in F#, the script for the generation of thermodynamically unfavored structures was written in Python, and all other analysis was done in MATLAB. The F# scripts and MATLAB scripts are available at: https://github.com/Gavin-J-Li/MLB-seq.

## Data Availability

All NGS raw data is available at: https://figshare.com/articles/dataset/Experimental_Confirmation_of_Multiple_Co-Existent_DNA_Secondary_Structures_using_Low-Yield_Bisulfite_Sequencing_Raw_Sequencing_Data_/14642214.

## Author contributions

JB, JL, BY, AP, and DYZ conceived the project. JB designed the oligo sequences and developed the experimental workflow. JB and JL performed the experiments. JL and BY aligned the NGS data to target oligos and performed statistical analysis. JL implemented the probability model. JL and BY validated the model. JL and MXW performed analysis on biological oligos. JL and DYZ wrote the manuscript with input from all authors.

## Additional information

DYZ is a co-founder and significant equity holder of Nuprobe Global, Torus Biosystems, and Pana Bio. JL was a consultant for Torus Biosystems during this research. AP and BY were fulltime employees of Microsoft during this research. MXW is a consultant for Nuprobe USA. There is a patent pending on the low-yield bisulfite sequencing.

