## Supplementary Material for "High-Throughput Measurement of Metastable DNA Secondary Structures using Multiplexed Low-Yield Bisulfite Sequencing (MLB-seq)"

### （Supplementary Information）

Jiaming Li, Jin Bae, Boyan Yordanov, Michael X. Wang, Andrew Phillips, and David Yu Zhang

(Dated: May 07, 2022)

### Supplementary Text

#### Detailed Experimental Protocol

(1) Amplifying Synthetic Oligos

In order to get enough amount of oligos, we assigned PCR primer binding regions and performed PCR. All oligo groups were divided into four larger groups so that each of them can be separately amplified. We normalized their concentrations for the final pooling. Primer pair 1 amplifies 372 oligos from groups 1-3, primer pair 2 amplifies 185 oligos from groups 1 and 3, primer pair 3 amplifies 355 oligos from groups 4 and 5, and primer pair 4 amplifies 1,057 biological oligos. Note that the 5' ends of all the 5' primers had 3 phosphorothioate modifications (denoted as '*' in 'LBY_Sequences.xlsx') to prevent any exonucleases activity, including Lambda exonuclease in step (3).

We first diluted the synthetic oligo pool by mixing 1 μL of 8 ng/μL oligo pool, 1 μL of yeast tRNA (1μg/μL), and 98 μL of TE buffer with 0.1% Tween-20 (Sigma-Aldrich). PCR mix was prepared by mixing 113.4 μL of nuclease-free water, 9.45 μL of DMSO (NEB), 157.5 μL of KAPA HiFi HotStart Uracil+ ReadyMix (KAPA Biosystems), and 3.15 μL of the diluted oligo pool. We then divided the mixture into four tubes, 67.5 μL each, and added 7.5 μL of one of the primer pair mixtures with 3 μM forward primer and 3 μM reverse primer. Next, we divided the mixture into each tube to three 25 μL aliquots and put them on the thermocycler for PCR. The PCR protocol was: 95 ºC for 5 min, 21 cycles of 98 ºC for 20 sec, 60 ºC for 15 sec, and 72 ºC for 1 min, and a final elongation stage at 72 ºC for 1 min. All mixing steps above were performed on ice.

We then mixed the three 25 μL aliquots back to 75 μL before the cleanup with 1.8x AMPure XP magnetic beads (Beckman Coulter), followed by the elution with 25 μL of TE buffer. After elution, all four tubes were quantified by Qubit, and were mixed to ensure that the final oligo concentrations from the four tubes was 8:1:8:24 in the final mixture for the optimal NGS uniformity. This mass ratio was determined empirically. Lastly, another 1.8x bead purification with AMPure XP was performed, followed by the elution with 16 μL of 0.1x TE buffer to concentrate the final oligo pool.

(2) Truncating the 3' Primer Regions with Restriction Endonuclease:

We truncated the 3' primer regions by mixing 9 μL of nuclease-free water, 3 μL of CutSmart Buffer (NEB), 15 μL of the oligo pool from last step, and 3 μL of BciVI restriction endonuclease (NEB), and incubating the mixture at 37 ºC for 2 hours and 80 ºC for 20 minutes. Then bead purification with 2.0x AMPure XP was performed, followed by the elution with 32 μL of 0.1x TE buffer.

(3) Digesting the Bottom Strand to create single-stranded DNA (ssDNA) and truncating the 5' primer region.

We used Lambda exonuclease (NEB) to digest the bottom strand of dsDNA. The phosphorothioate modification on the 5' end of the top strand and the exposed phosphate group on the 5' end of the bottom strand after BciVI digestion ensured that only the bottom strand was specifically digested. The degradation was then followed by USER Enzyme (NEB) treatment to truncate the 5' primer region.

The mixture was 11 μL of nuclease-free water, 3 μL of CutSmart Buffer, 15 μL of oligo mix from last step, and 1 μL of Lambda exonuclease. The mixture was incubated at 37 ºC for 30 minutes, 75 ºC for 10 minutes, and maintained at 4 ºC. Next, 1 μL of USER Enzyme was added with the tube remaining on the thermocycler, and the mixture was then incubated at 37 ºC for 30 minutes. The final oligo mix was purified with the Oligo Clean & Concentrator Kit (Zymo Research) and eluted with 1x TE buffer.

(4) Pre-annealing and Bisulfite Conversion

Before sodium bisulfite treatment, we pre-annealed the final oligo mix after adding 5 M NaCl 20% v/v. The mixture was heated at 95 ºC for 3 minutes, followed by cooling with the rate of -1 ºC/10 s down to 25 ºC and incubation for 3 minutes.

To prepare the sodium bisulfite solution, we first degassed the nuclease-free water with ultrasonication for 10 minutes. We dissolved 950 mg of sodium metabisulfite (Sigma-Aldrich) with 9.5 mL degassed water by gently shaking. We then added 80 μL of 10 M NaOH (Sigma-aldrich), mixed by gently shaking, and added 120 μL of 250 mM Hydroquinone solution (Acros Organics, 99%). Next, we carefully added 10 M NaOH until the pH of the solution reached 5.5. Finally, we added water until the volume reached 10 mL. It is critical to use freshly prepared sodium bisulfite solution.

For bisulfite conversion, we mixed 6 μL of pre-annealed oligo mix and 114 μL of the sodium bisulfite solution, and incubated the mixture at 55 ºC for 0, 2, 4, or 6 hours. Before performing cleanup with EZ DNA Methylation-Gold Kit (Zymo Research), we added 5 M NaCl to the binding buffer in order to make it have 0.833 M NaCl for salinity compensation. Other steps followed the protocol of the kit. All sample were eluted with 11 μL of elution buffer and stored at -20 ºC until the next step.

(5) 3'-Ligation of Illumina P7 Adapter

To further proceed to NGS, we first performed ligation to attach Illumina P7 adapters to the 3'-ends of the oligos, following the SPLAT method by Amanda Raine, et. Al [28]. We prepared the ss1 mix by making 40 μM LB-splint1 and 60 μM LB-adap3 in 49 μL TE buffer and added 1 μL of 5 M NaCl. Incubation started from 65 ºC for 1 minute, followed by cooling at the speed of -1 ºC/10 s down to 25 ºC. The annealed ss1 mix was stored at 4 ºC before use.

For the 3' ligation process, we mixed 2.5 μL of nuclease-free water, 2.5 μL of 40% PEG 4000 (Sigma-aldrich, re-made every 3 months), 2 μL of T4 DNA Ligation Buffer (NEB), 10 μL of the sample from the last step, 2 μL of ss1 mix, and 1 μL of T4 DNA Ligase (NEB). The mixture was incubated at 20 ºC for 1 hour and 65 ºC for 10 minutes.

After incubation, gel extraction was performed to size-select the ligated oligos. We ran the oligo sample through gel electrophoresis with 2% agarose gel to separate the oligos by size and used the Zymoclean Gel DNA Recovery Kit to extract the oligos we want. We cut the gel covering the two bands right above the 100 nt marker for the recovery and eluted with 11 μL elution buffer.

(6) 5'-Ligation of Illumina P5 Adapter

We then attached Illumina P5 adapters to the oligos with the second ligation. The protocol was identical to 3'-ligation in step (5), except that ss2 mix (prepared in the same way as ss1 mix but using LB-splint2 and LB-adap5) was used in place of ss1 mix. The reaction was followed by bead purification with 6x AMPure XP at and elution with 15 μL of 0.1x TE.

(7) Quantification, Amplification, and NGS

For quantification before the final PCR, we performed qPCR to determine the appropriate cycles for the amplification. KAPA HiFi HotStart Uracil+ ReadyMix was used according to the manual except for adding 3.3% v/v DMSO and 3x SYBR Green (Thermo Fisher). NEBNext Multiplex Oligos for Illumina Set 1 (NEB #E7600) was used as the primers. For PCR, we used the same protocol as qPCR except SYBR Green. Finally, we performed bead purification twice with 1.2x beads. The first and the second purification used 26 μL and 15 μL of TE for the elution, respectively.

**Libraries and Experimental Variables**

In the experiment, we produced 6 libraries all with the same 1,969 input oligos but with varied experimental conditions. We had two variables: the DNA concentration during the bisulfite reaction and the 55ºC incubation time of the reaction (in step (4) of Detailed Experimental Protocol). The specific varied conditions are shown in Table S1. We used data from library 1 for all analysis and demonstration in the main text and supplementary materials, if not specially specified.

The NGS read depth of each library is plotted in Fig. S2. In the end, the majority of oligos had sufficient NGS read depths across all libraries, demonstrating the preliminary robustness of the experimental workflow.

The CY at varied experimental conditions are compared in Fig. S3. We plotted the CY of each C nucleotide of one group 1 oligo through different libraries. The CY increased as the reaction hours got longer but did not show any correlation with the oligo concentration just identifying by our libraries.

To test the reproducibility of MLB-seq, we compared the CY of each same C loci between different libraries (Fig. S4). The CY of each C nucleotide in one library is plotted on x-axis, and the CY of the same loci in another library is plotted on y-axis. Each plot covers all the C nucleotides from all 1,969 oligos. The linear fitting results and the correlation coefficient $R$ are shown. Note that the x- and y-axis scales varied because the overall conversion yield varied among different libraries due to varied experimental conditions. We found very good reproducibility among libraries 1, 4, 5, 6, with R close to 1. Library 3 was an outlier because the overall conversion yield was too low.

**Conversion Yield of Dangle, Loop, and Stem of Each Library.**

We summarized the different CY of C nucleotides at dangle, loop, and stem regions of all libraries in Fig. S5. Excluding Library 2 which had an incubation time of 0 hour, as the negative control, the data from all other libraries showed significant discrimination in the CY of open and closed C nucleotides.

**Conversion Yield of C Nucleotides at Mismatches**

Here we plotted the CY of C nucleotides at the mismatches, next to the CY of C nucleotides at dangle, loop and stem regions for comparison (Fig. S6). Overall, C nucleotides at mismatches had a mean CY of 9.9%, which is between that of C nucleotides at dangle and loop and significantly higher than that of C nucleotides at stem regions.

When looking at the histogram, the CY of C nucleotides at mismatch positions had a wide distribution starting from 0% to higher, overlapping with that of C nucleotides both at stem regions and at loop and dangle regions, though. We speculate that this is because the mismatched positions were designed to have sizes of only 1, 2, or 3 nt. The C nucleotides on these mismatched positions, even though open by themselves, are right next to, and thus constrained by, the duplex stem nucleotides. Therefore, when the neighboring duplex stem is stable (e.g. when they are dense with C and G nucleotides), the mismatched C nucleotide might have a rather low chemical accessibility as if it is closed. On the other hand, when the neighboring duplex stem is not stable and allows for chances of base breathing at the end of the stem, then the mismatched C nucleotide might bear less constraint and have a higher chemical accessibility that resembles an open C nucleotide.

**Conversion Yield of C Nucleotides at Hairpin Loops with Varied Loop Lengths**

The full plot of the CY of C nucleotides at hairpin loops with varied loop lengths is shown in Fig. S7, as a supplement to Fig. 2b. As the loop length grew longer, the effect it had on the CY of C nucleotides in the loop became less and neglectable.

**Conversion Yield of C Nucleotides of oligos in Groups 2 and 4**

We analyzed the CY of C nucleotides at the loop, dangle, and stem regions with oligo targets from Groups 2 and 4 (Fig. S8a, b). We found the distinction between open and closed C nucleotides by their CY were consistently significant for these oligos. However, it is noticeable that there were a minority of loop C nucleotides among Group 4 oligos with very low CY, which overlapped with the stem C nucleotides (as marked by the brown arrowhead in Fig. S8b). Inferred from our findings in Fig. 2c, d, we hypothesized that some of these loop C nucleotides with low CY might have resulted from them being next to the duplex stem. Since all Group 4 oligos were each designed to have multiple hairpins, there were more chances for loop C nucleotides to be right next to the duplex stem region. To validate this hypothesis, we excluded the CY of all C nucleotides at the joint between loop and stem, and between dangle and stem (Fig. S8c) and replotted the histograms. Now the overlapped CY of loop and stem C nucleotides near 0% did decrease significantly (Fig. S8d).

**Calculation of the Posterior Conditional Probabilities**

To develop the model for calculating the probability of each NGS read corresponding to each candidate structure given its original sequence and conversion data, we first need to determine the posterior conditional probabilities of a C nucleotide being open or closed given it was observed converted or unconverted: *P(SO|BC)*, *P(SC|BC)*, *P(SO|BU)*, and *P(SC|BU)*, where *BC* and *BU* stands for bisulfite converted and unconverted, and *SO* and *SC* stands for structure open and closed, respectively. We derived these posterior probabilities from experimental conversion data of the positive control oligos. Note that the conversion yield varied with different libraries, so we should derive the probabilities from conversion data of positive control oligos that underwent the identical experimental conditions as the target oligos that we plan to study with the model.

We counted and summarized the C nucleotides that were (1) open and converted, (2) open and unconverted, (3) closed and converted, and (4) closed and unconverted from all oligos in Groups 1, 2, and 4, as shown in the Table S2. Then, we calculated the four posterior probabilities following equations below, where N stands for the summed number of C nucleotides in the last row of Table S2.

$$P(SO|BC) = \frac{N_{SO\&BC}}{N_{SO\&BC} + N_{SC\&BC}}$$

$$P(SC|BC) = \frac{N_{SC\&BC}}{N_{SO\&BC} + N_{SC\&BC}}$$

$$P(SO|BU) = \frac{N_{SO\&BU}}{N_{SO\&BU} + N_{SC\&BU}}$$

$$P(SC|BU) = \frac{N_{SC\&BU}}{N_{SO\&BU} + N_{SC\&BU}}$$

In our specific case with conversion data from Library 1, we finally got the four conditional probabilities: $P(SO|BC) = 0.93$, $P(SC|BC) = 0.07$, $P(SC|BU) = 0.45$, $P(SO|BU) = 0.55$.

**Internal Variables of the Model**

There are two important internal variables that should be addressed in the model as defined below:

(1) *Tolerance,* denoted as $t$. We use *tolerance* to classify whether an NGS read is non-ambiguous or non-specific.

We denote the log-scaled probabilities of the read corresponding to structure 1 and structure 2 as $p1$ and $p2$ respectively. If $abs(p1 - p2) < t$, then this read is consistent with both structures and classified as non-specific. Otherwise, this read is classified as non-ambiguous and the structure with higher probability wins.

(2) *C Number*, denoted as $n$, the number of C nucleotides in the oligo target.

These two variables are important because, theoretically, a *tolerance* too low might result in false positive or negative and reduce the specificity, and a *tolerance* too high might classify too many reads as non-specific and limit the sensitivity. All figures in the main text and in the Supplementary are processed when *tolerance* is set at 0.1, if not specifically noted. We also need to investigate whether the number of C nucleotides in the target sequence affects the model performance, since MLB-seq only probes C nucleotides.

**Performance Test on the Internal Variables via Simulation**

Via thorough performance test, we determined the optimal value for *tolerance*, and we validated that the model performs stably well despite the varied *C Number*.

We first used generated dataset via simulation as the model input. Compared to experimental MLB-seq data from positive control oligos, simulation generates excessively larger data size at minimal time and cost and thus can provide us with data covering a wide range of variations.

We ran the simulation following the steps below:

(1) Randomly generate an oligo at 100-nucleotide length with A, T, C, G bases all at 25% probability (not actual fraction).
(2) Use Nupack to calculate the MFE structure and extract the binary open/closed pattern of all C nucleotides. This pattern will be used as the positive reference.
(3) Randomly generate a binary open/closed pattern, whose length is the same as *C number* of the oligo target. This pattern will be used as the negative reference. If the negative reference is identical to the positive reference, then generate another different pattern to replace the existing one.
(4) Generate an array of zeros at the length of *C number*. Go through the array and for each element, randomly convert it to 1 at the probability of $P(BC|SO)$ if it corresponds to an open C in the MFE structure, and at probability of $(BC|SC)$ if it corresponds to a closed C in the MFE structure. This mimics the conversion data from one NGS read. Repeat this step for a large dataset of this oligo target.
(5) Feed the data from step (4) and the open/closed patterns from steps (2) and (3) into the model and obtain the output matrix of log-scaled probabilities. Screen out the non-specific reads using *tolerance*. Then calculate the supporting rates for the two structures using only the non-ambiguous reads.
(6) repeat steps (1) to (5) for a different oligo target.

We varied the *tolerance* value from 0 to 2 and generated 50 random oligo targets at each datapoint. For each oligo target, we generated 10,000 reads as the conversion data. Fig. S10a summarizes the supporting rates for the Nupack MFE Structures, i.e. the true positive rates, from all simulated oligo targets. Fig. S10b summarizes the fraction of non-ambiguous reads. We found that when the *tolerance* is set too small (≤ 0.1), there are significant false negative supporting rates. When the *tolerance* goes larger (≥ 0.2), the fraction of non-ambiguous reads was reduced, resulting in lower sensitivity. Thus 0.1-0.2 would be the optimal value for *tolerance*.

We then hard-set the number of C nucleotides in the sequence in step (1) of the simulation, and acquired the supporting rates for the Nupack MFE structures and the fraction of non-ambiguous reads at varied *C numbers*, as summarized in Fig. 9c, d. Here the *tolerance* is set at 0.2. Looking at both the supporting rates and the non-ambiguous reads, the model performs robustly with *C number* ranging from 5 to 100.

**Performance Test on the Internal Variables Using MLB-seq data from Positive Control Oligos**

We did further performance test with experimental data from all positive control oligos in Groups 1, 2, and 4 and validated the optimal *tolerance* value and the model robustness against varied *C number*.

We varied the *tolerance* values at 0.1, 0.2, and 0.3, and processed the MLB-seq data from all positive control oligos. The supporting rates for the Nupack MFE structures, i.e. true positive rates, are summarized in Fig. S11a, b, c. We found that when *tolerance* = 0.1, the true positive rates are high for almost all positive control oligos. As the *tolerance* increases, there are some oligos that give ~ 0% supporting rates for the Nupack MFE structures. We further looked into the model output probability matrix of these oligos and found that these are all false negatives. Fig. S11d shows an example. Almost all NGS reads have a higher probability corresponding to structure 1, i.e. the Nupack MFE structure, but these reads are all classified as non-specific reads because the *tolerance* value is larger than the probability difference between two structures. Thus, we can conclude that a *tolerance* too large will result in an increase of false negatives.

We also summarized the fraction of non-ambiguous reads at varied *tolerance* values (Fig. S11d). When $tolerance = 0.3$, there is a significant number of oligos whose majority of reads are classified as non-specific. This is consistent with the false negatives from the supporting rates. According to both the supporting rates and the non-ambiguous reads, we conclude the optimal value for *tolerance* is 0.1.

The *C numbers* of these positive control oligos vary from 24 to 44. We plotted the supporting rates for Nupack MFE structures against *C numbers* in Fig. 4e and found no correlation between the two. Combining results from previous simulation with a much wider *C number* range, we conclude that the model performs robustly despite the variation of *C number*.

**Generation of Thermodynamically Unfavored Structures**

We are seeking to generate diverse structures to validate our algorithm, but random sampling generally results in generating unfeasible structures with high energies. Enumerating sub-optimal structures, on the other hand, produces structures that are energetically favourable but are generally similar to the mfe structure.

To balance these trade-offs, we treated the task of generating diverse and energetically feasible (though still thermodynamically unfavoured compared to the mfe) structures, as a multi-objective optimization problem. As one objective, we seek stable structures (minimum energy) and as another objective we require that these structures be different from the known mfe (structure distance maximization).

To implement this strategy, we adapted ideas from multi-objective evolutionary algorithms [29], where we focus our search on non-dominated (Pareto-optimal) structures - structures that are either lower energy or further from (less like) the mfe compared to all other ones. We start with a population of randomly generated, valid structures (e.g., where only standard base pairs and no pseudoknots are allowed). At each iteration, we randomise the set of structures with operations commonly used within genetic algorithms (mutation and cross-over), while we preserve only the resulting structures that are valid but also non-dominated. The set of structures that we consider for the evaluation of the algorithms is produced after several iteration of this optimization. We found that the Pareto front got solidly improved as the iteration moved forward (Fig. S12). In the end for each oligo, we have a pool of structures with free energy ranging from mfe to slightly above 0 kcal/mol, which can serve as stable candidate structures (with negative free energies) or thermodynamically unfavored negative-control structures (with positive free energies).

For the validation of the analytical model (Fig. 4 a-d), we randomly selected a structure as the thermodynamically unfavored comparison structure following the two restrictions below: (1) the structure has at least 10 bases at different open and closed states from the mfe structure; (2) the free energy of the structure is above -2 kcal/mol.

**Information of the Biological Sequences**

We selected 1,027 biological sequences from 212 human genes curated at COSMIC following the steps below: (1) downloaded the 212 curated genes from COSMIC (their alignment locations on the chromosomes are shown in Fig. S6a); (2) Calculated the partition function ($\Delta G$) of multiple 100-nt fragments of each gene (e.g. nt 1-100, 101-200, 201-300, etc. ); (3) Remove the fragments with >6 nt homopolymer stretch; (4) select 4 fragments with the lowest $\Delta G$ and 1 or 2 fragments with medium $\Delta G$ from each gene. These steps helped ensure that the 1,027 biological sequences selected have a wide distribution of GC contents and $\Delta G$ (Fig. S13c). We later calculated their mfe using Nupack and confirmed that their mfe also cover a wide range (Fig. S13b). In the end, 193 of the 1,027 oligos are truncated from exon, 864 oligos are from intron, and 58 oligos span through intron and exon regions.

**Necessity of Introducing the Pairwise Comparison and Condorcet Winning Strategy**

When an oligo target has multiple candidate structures within the 2 kcal/mol energy gap predicted by Nupack, the log-scaled probability matrix directly output from the model that compares all structures together might generate too many non-specific reads (reads that supports ≥2 different structures) and make it hard further analysis including identifying the dominant structures and calculating the supporting rates. To deal with this, we made pair-wise comparisons of the log-scaled probabilities between each pair of structures and selected the top three winning structures for a final comparison (Fig. S14b). Thus, our final output was narrowed down to just these three winning structures so that a sufficient fraction of the NGS reads could now give us specific information on structure distributions (Fig. S14c).

**Conversion Yields of Open and Closed C nucleotides from Biological Sequences with Two or More Dominant Structures**

We present the histograms of CY of all open and closed C nucleotides calculated using only one or several subsets of NGS reads R1, R2, and R3. All the histograms from the 'correct' combinations, i.e. the first two in Fig. S15a and the first and third in Fig. S15b, show distributions of open and closed C CY more consistent with our previous observation than those from the other 'incorrect' combinations.

**Fig. S1.**
Detailed experimental workflow diagram.

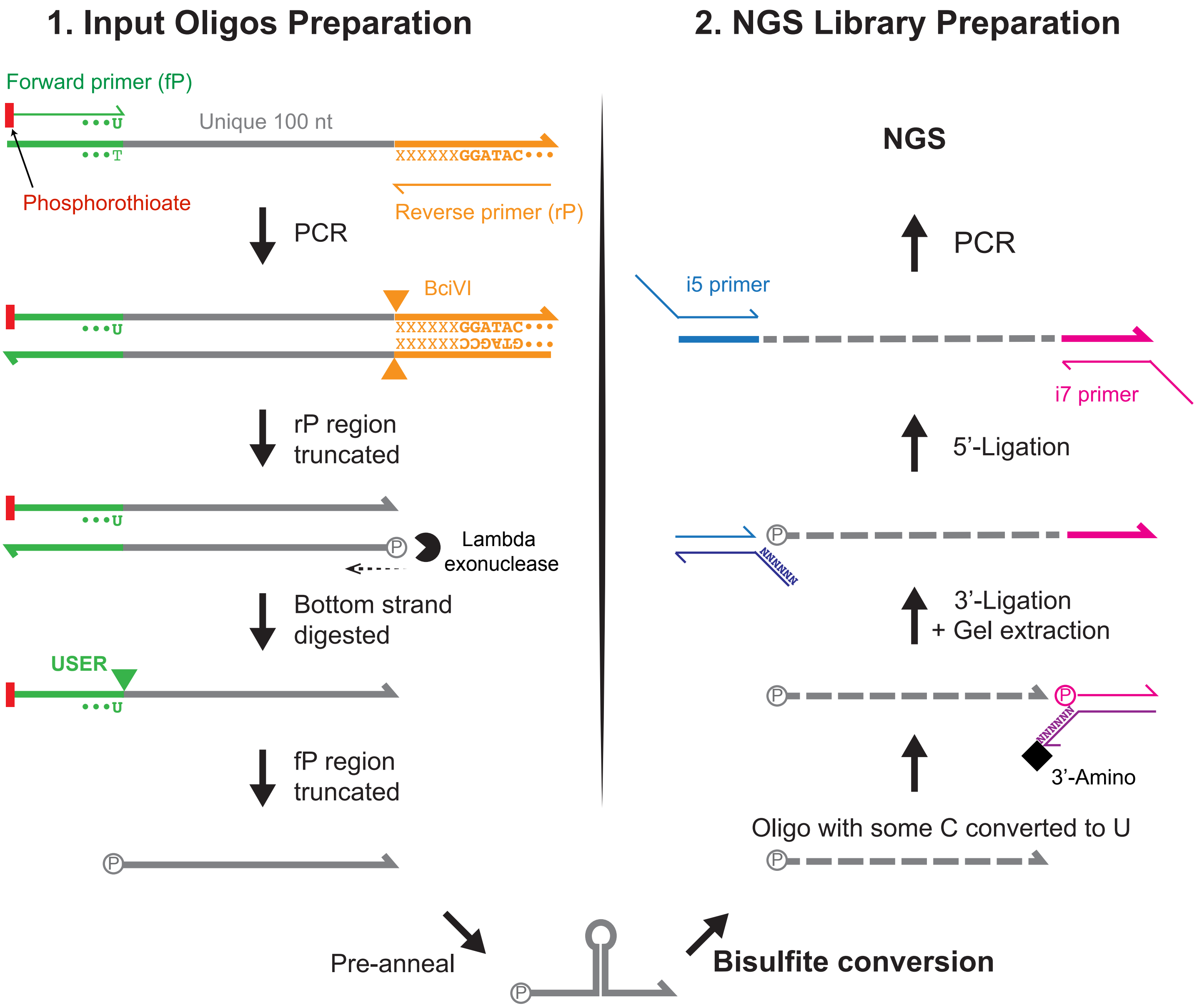

**Fig. S2.**
The NGS read depth of each library.

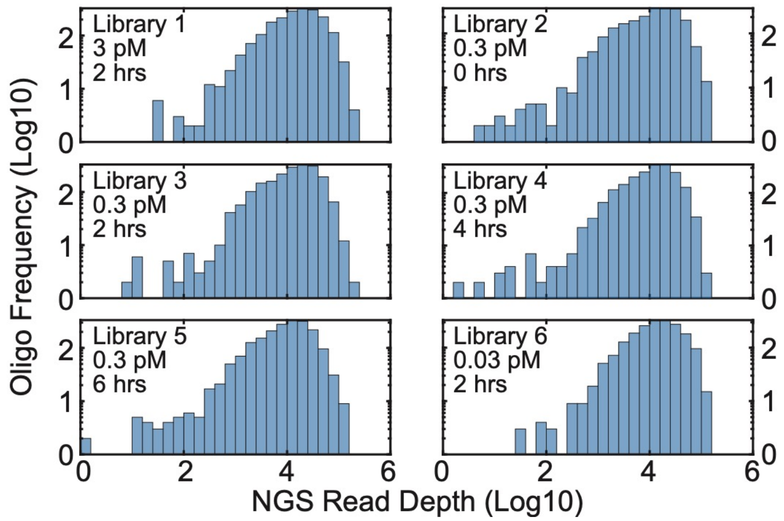

**Fig. S3.**
The change of overall CY at varied experimental conditions.

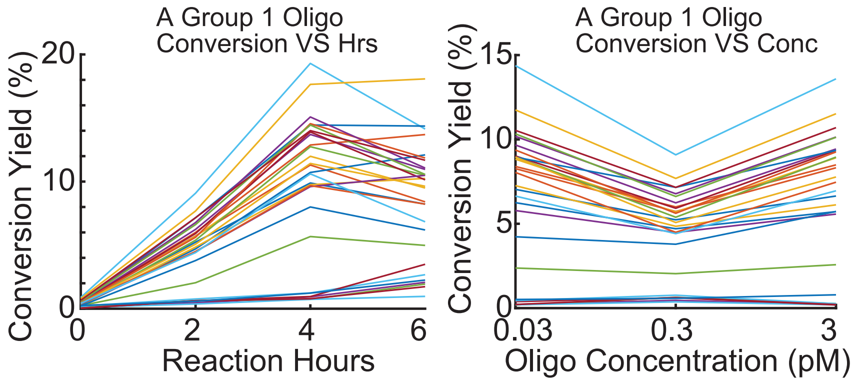

**Fig. S4.**
Reproducibility of all 6 libraries. For each of these plots, the CY of all C nucleotides from one library are plotted on the x-axis, while the CY of the same C nucleotides from another library are plotted on the y-axis. Note that all ~58,000 C nucleotides from all 1,969 oligo targets were taken into account and their CY were utilized for the calculation of linear fitting and correlation coefficient R. But for easier viewing, only the CY of 5,000 randomly selected C nucleotides were plotted in the graphs.

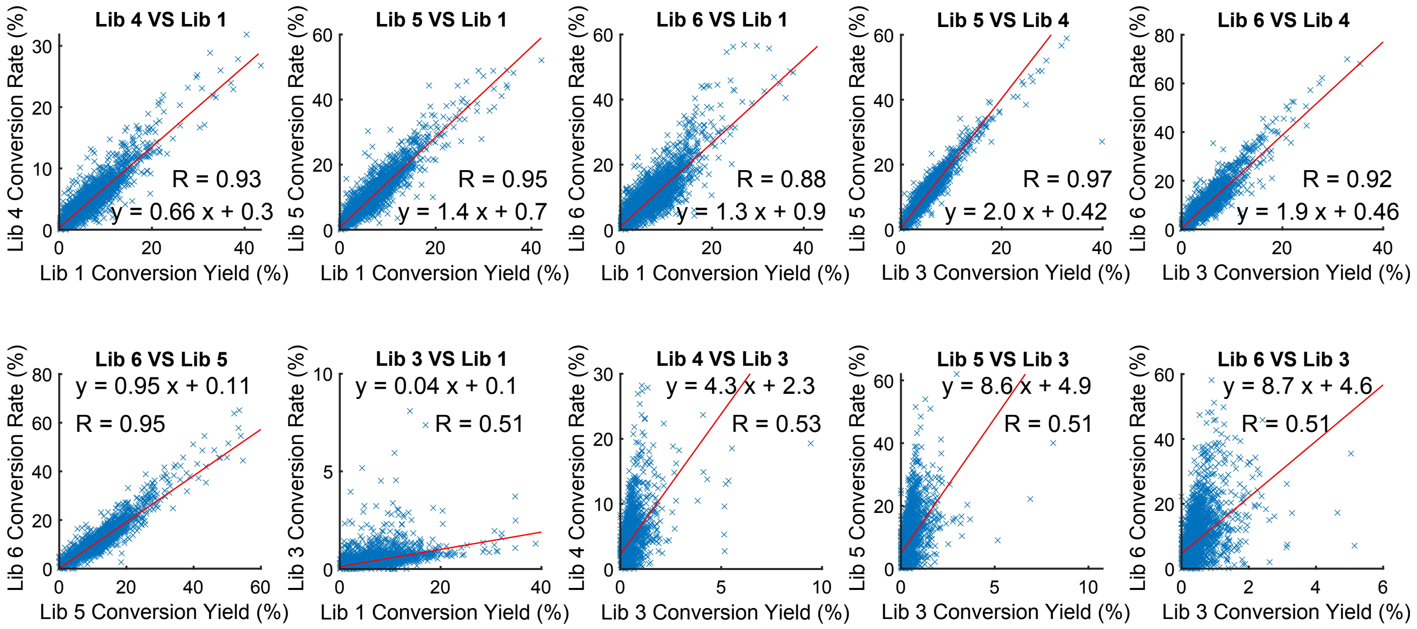

**Fig. S5.**
Differentiated CY of dangle, loop, and stem of each library. The mean values of each histogram are marked by the red dashed lines.

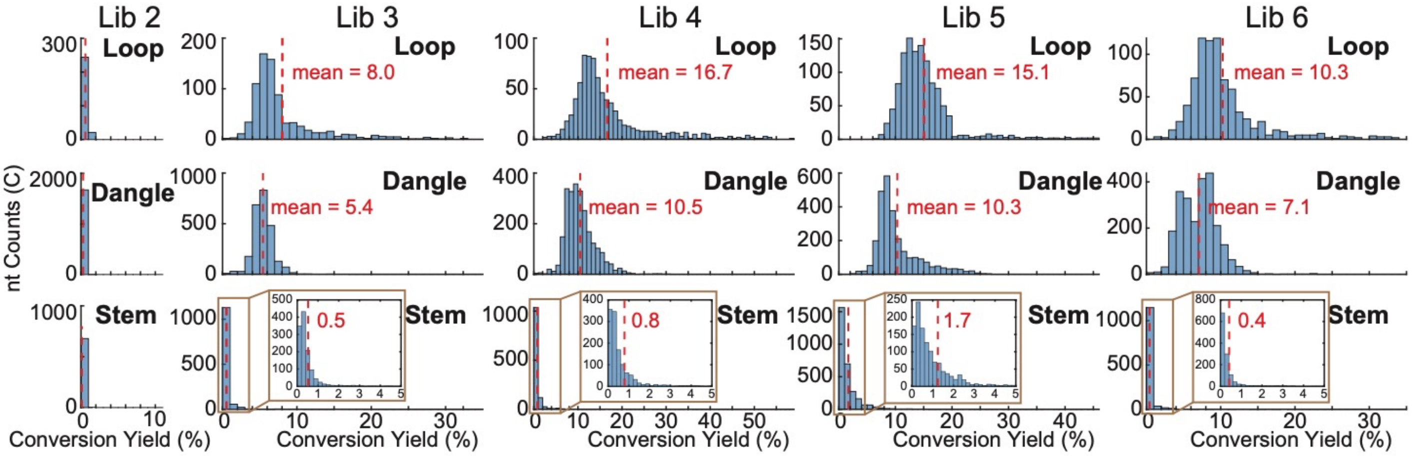

**Fig. S6.**
CY of C nucleotides at mismatched positions, compared to that of C nucleotides at loop, dangle, and stem regions.

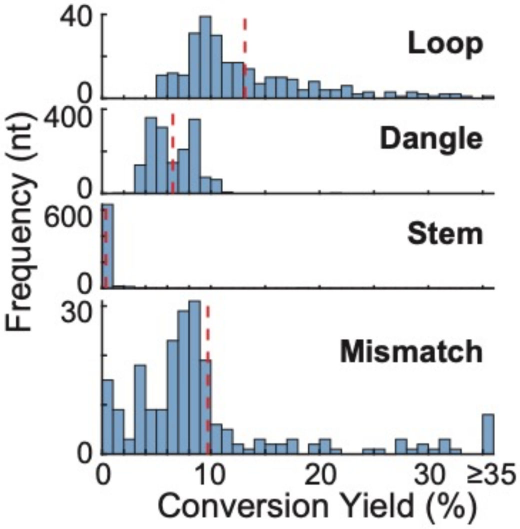

**Fig. S7.**
Conversion yields of C nucleotides at hairpin loops with varied loop lengths.

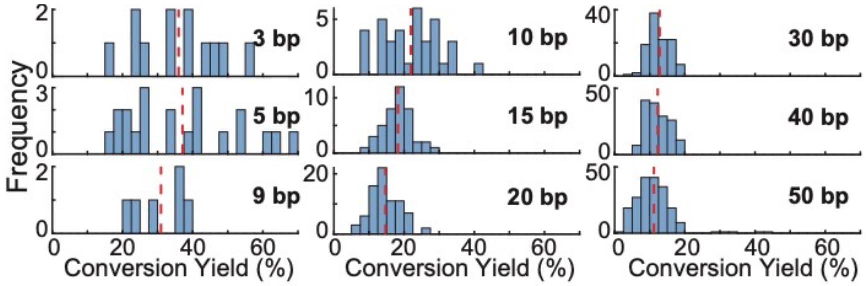

**Fig. S8.**
CY of C nucleotides at loop, dangle, and stem regions from Group 2 and 4 oligos.

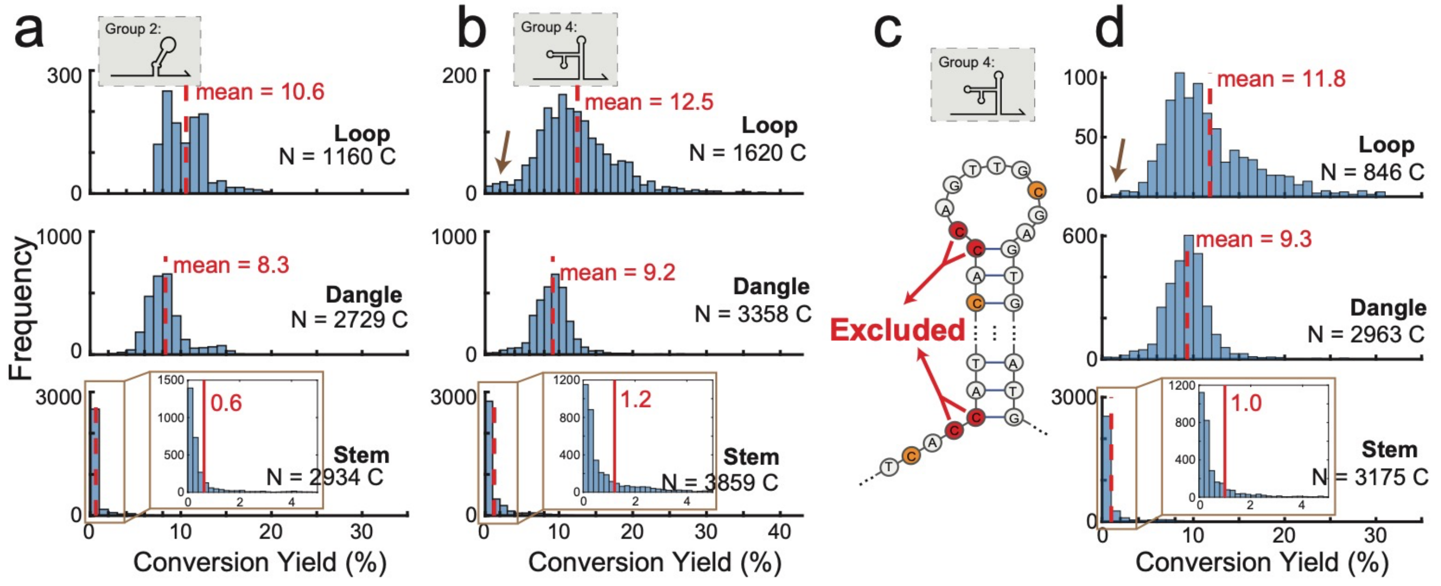

**Fig. S9.**

Plot of CY from MLB-seq against the Nupack predicted base open probabilities (OP). The CY was calculated from MLB-seq data. The OP was predicted by Nupack 'pairs' function. The colorbar represents the density of C nucleotides on each plotted spot. 'Total N' represents the number of C nucleotides plotted in each insertion. The red lines are linear fitting using all data. 'R' represents the correlation coefficient of linear fitting.

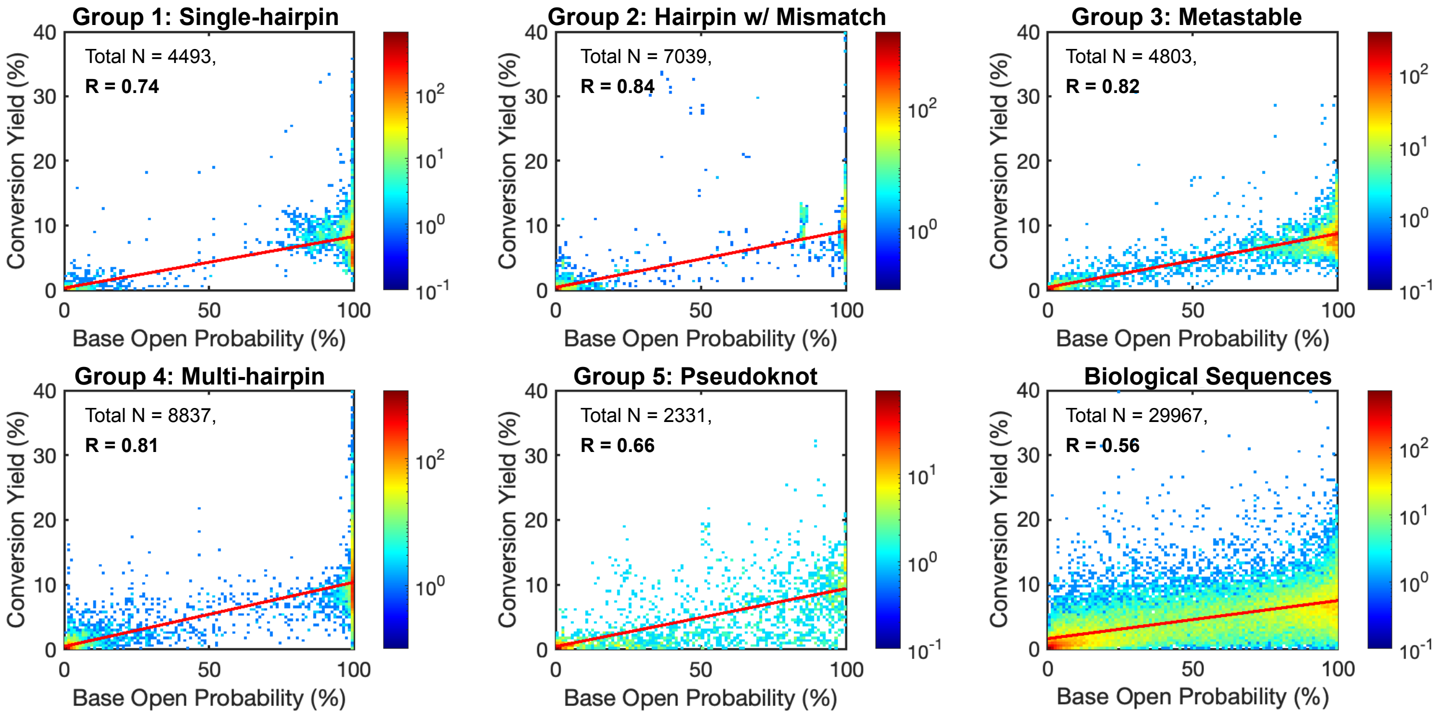

**Fig. S10.**
Performance test on varied tolerance and C number via simulation.

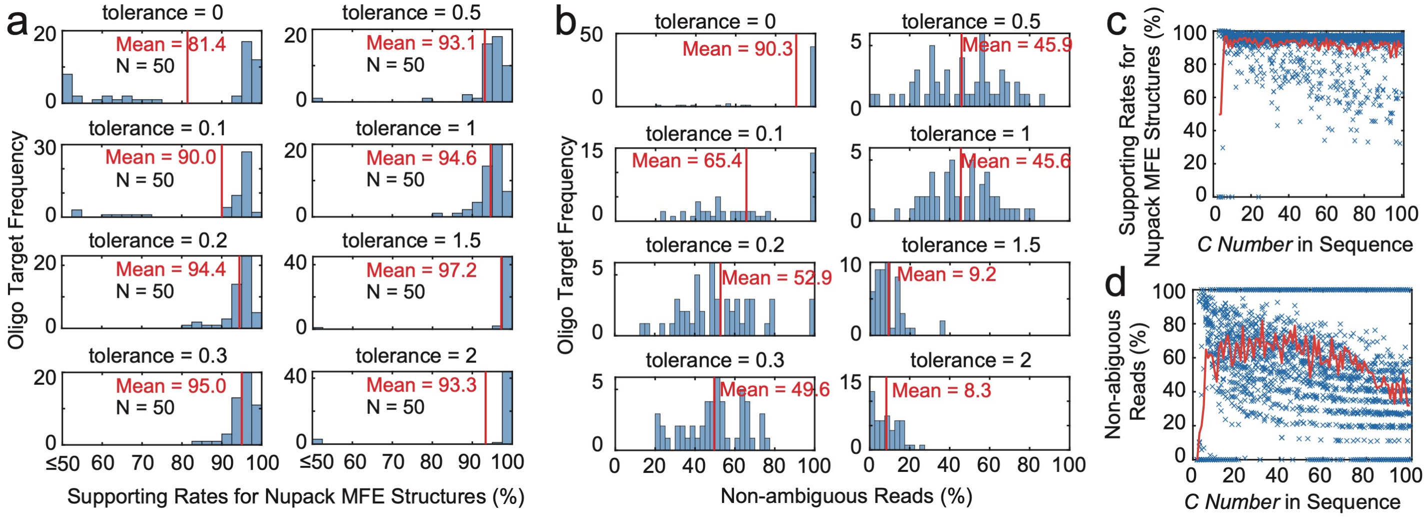

**Fig. S11.**

Performance test on varied tolerance and C number using MLB-seq data from positive control oligos

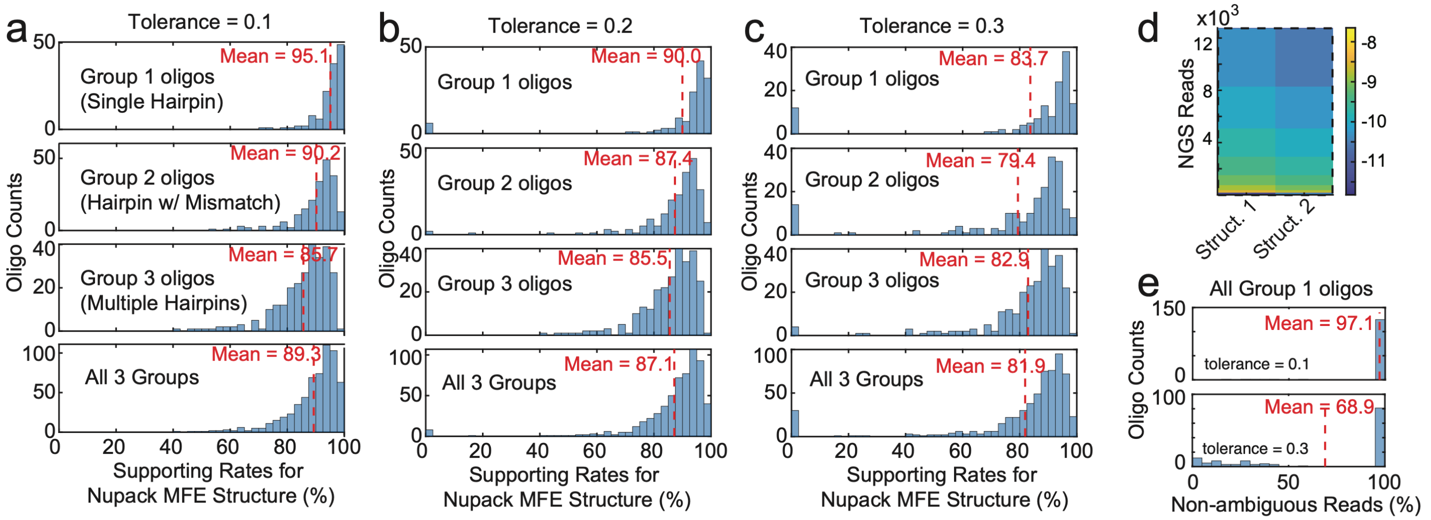

**Fig. S12.**

The iteration traces of the Pareto front being optimized. (a) from an example oligo target from Group 1; (b) from an example oligo target from Group 4. Each datapoint represents a structure preserved after each iteration. With the same distance from the mfe structure (same y-axis value), all datapoints steadily moved to the left, showing that their free energies got reduced as iterations moved forward.

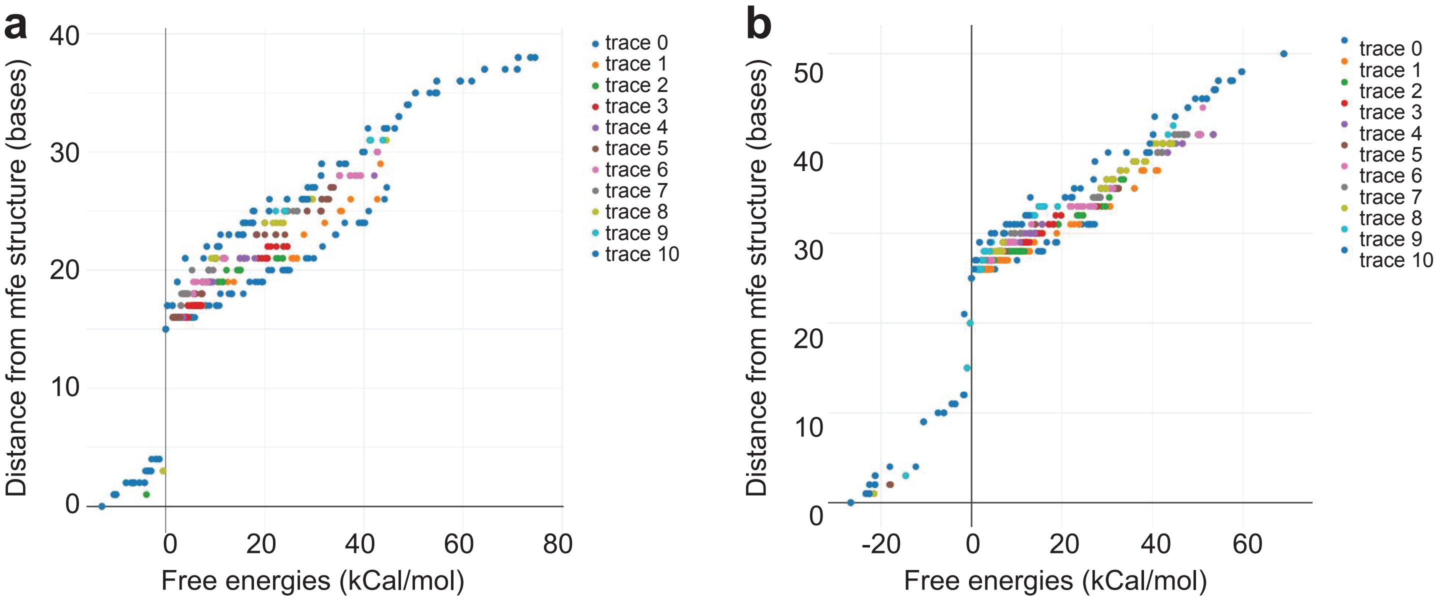

**Fig. S13.**
Information of the 1,027 biological sequences. (a) chromosome position mapping of the 212 gene that these sequences were truncated from. (b) mfe of these sequences calculated by Nupack. (c) GC contents of these oligos.

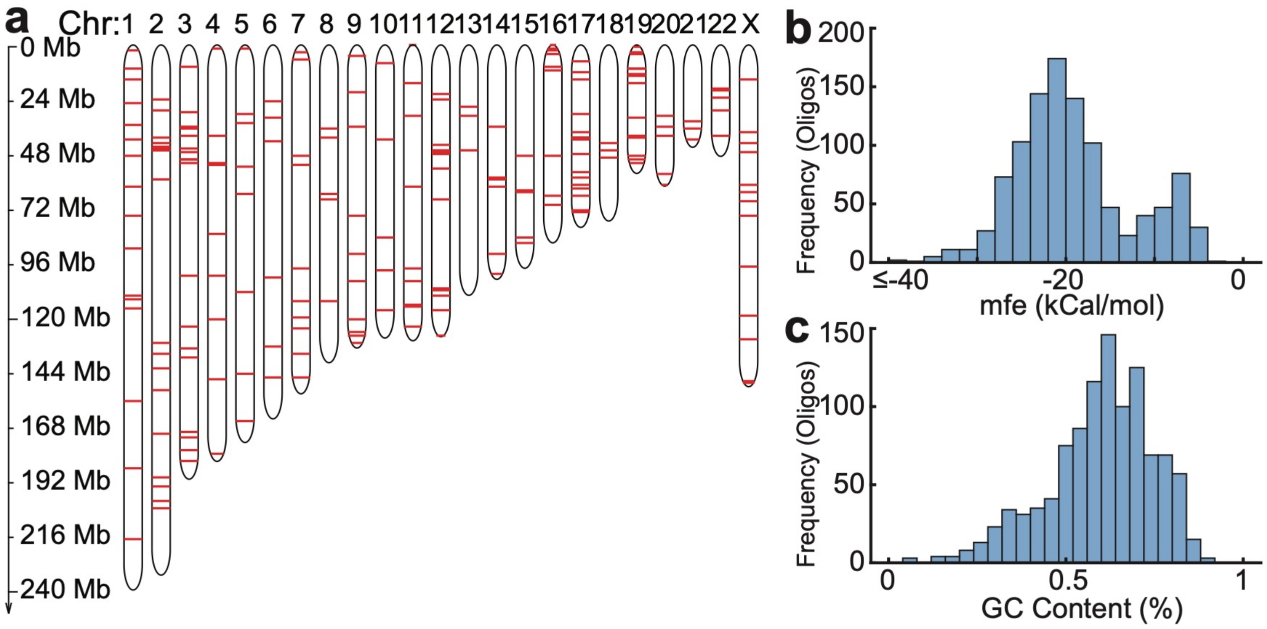

**Fig. S14.**
(a) Heatmap of the log-likelihood output matrix from the model, with 5 candidate structures. (b) Pairwise comparison winning strategy assigned structure 5, 4, and 2 as the three winning structures for final output. (c) Heatmap of the log-likelihood of all NGS reads of this oligo target corresponding to the three winning structures.

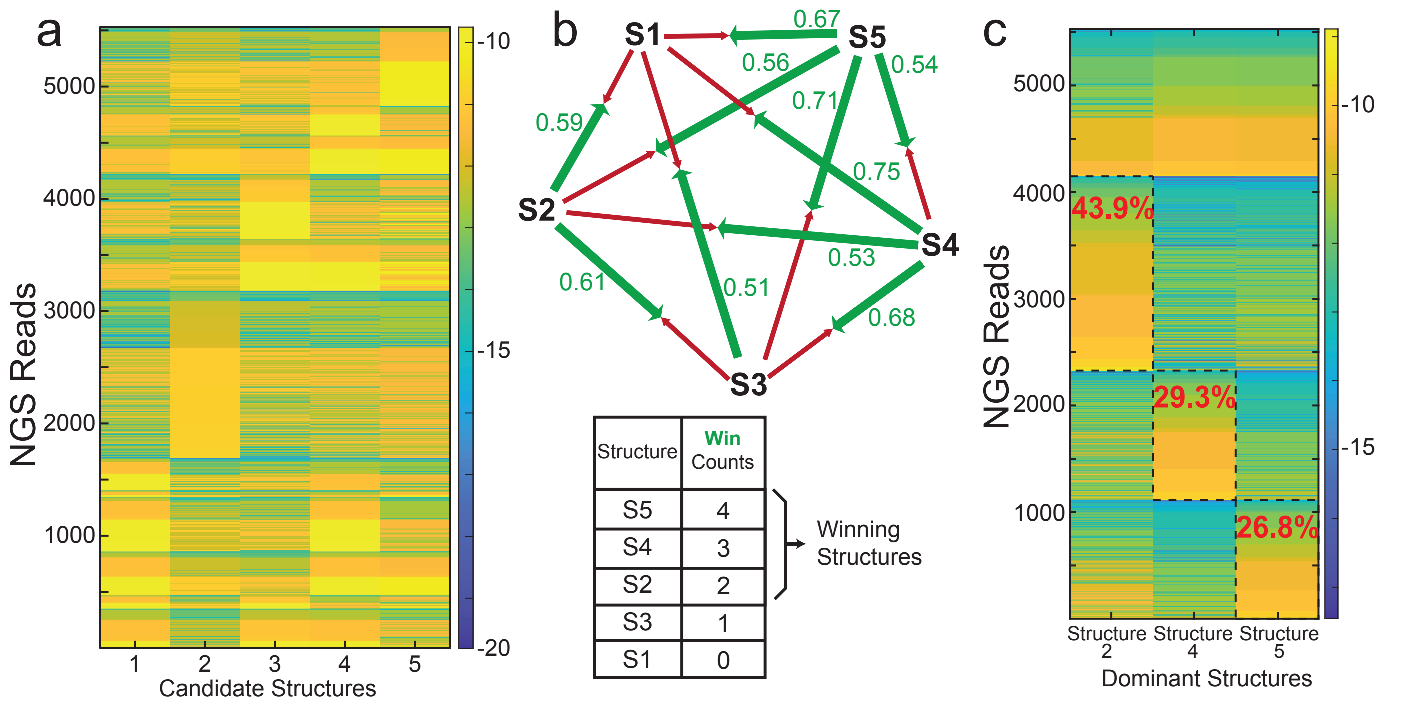

**Fig. S15.**
Evaluation of the model output on biological oligos with (a) 2 or (b) ⩾ 3 structures. In the 'Correlations' diagrams, the shaded orange represented the supposedly correct correlations between reads and structures supported by our model. The red box represented the reads and structure(s) we used calculate the CY and define the open and closed states. The CY for all open and closed C nucleotides are plotted in the histograms respectively.

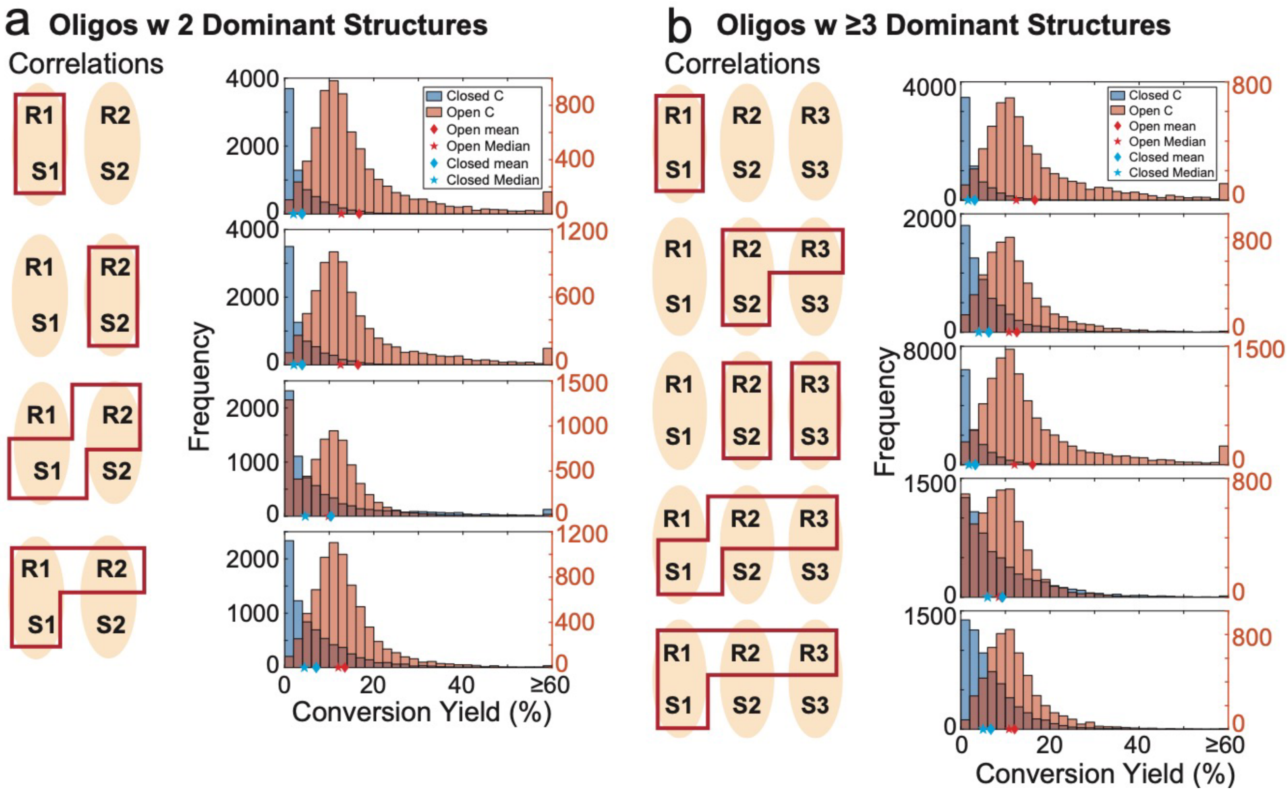

**Table S1.**
Experimental variables of the 6 NGS libraries.

| Library # | Oligo Concentration (pM per oligo) | Incubation Time (Hours) |
|---|---|---|
| 1 | 3 | 2 |
| 2 | 0.3 | 0 |
| 3 | 0.3 | 2 |
| 4 | 0.3 | 4 |
| 5 | 0.3 | 6 |
| 6 | 0.03 | 2 |

**Table S2.**

Number of C nucleotides that were (1) open and converted, (2) open and unconverted, (3) closed and converted, and (4) closed and unconverted summed from all positive control oligos.

| | SO & BC | SO & BU | SC & BC | SC & BU |
|---|---|---|---|---|
| Group 1 oligos | 4,581,517 | 51,583,397 | 78,085 | 19,641,223 |
| Group 2 oligos | 14,804,940 | 139,797,184 | 769,821 | 131,025,620 |
| Group 4 oligos | 43,024,593 | 397,421,931 | 3,730,326 | 321,771,559 |
| Sum of all above | 62,411,050 | 588,802,512 | 4,578,232 | 472,438,402 |

**Data S1. LYB_Sequences.xlsx (separate file)**
Sequences for all DNA oligos used in the experiments.
